# Functional ultrasound imaging in free-moving marmosets reveals that intra-call prosodic features modulate large-scale cortical dynamics

**DOI:** 10.64898/2026.11.02.763591

**Authors:** Xiaochun Cheng, Yichao Luo, Bingqing Wang, Mengdi Cao, Siyi Wu, Binshi Bo, Kaiwei Zhang, Mengchao Pei, Liwen Wan, Zhifeng Liang

## Abstract

Vocal turn-taking relies on sequential structure and rhythmic timing, yet how primate brains integrate these features across timescales in naturalistic, free-moving contexts is unclear. We developed functional ultrasound imaging for free-moving marmosets, combining an exogenous marker-based workflow for robust registration from non-standard imaging planes with preprocessing pipeline optimized to reduce locomotion artifacts. We systematically perturbed sequential and rhythmic features of interactive phee calls across hierarchical timescales. Cortical networks showed selective sensitivity to intra-call structure: disrupting phrase sequences increased activity in auditory and para-insular cortices and strengthened inter-subject functional connectivity within this network. Rhythmic manipulations primarily affected core auditory cortex: shortening inter-phrase intervals enhanced responses, whereas lengthening intervals suppressed them. These findings establish a practical framework for free-moving primate neuroimaging and identify intra-call prosodic features as promising targets for mechanistic investigation.

## Introduction

Vocal communication lies at the heart of social interaction, and turn-taking is among its most efficient organizing principles: partners alternate signals with short, structured gaps that sustain a stable exchange. Although frequently viewed as a distinctive feature of human conversation, turn-taking is not a behavior unique to humans.^1–4^ The common marmoset is particularly well suited for mechanistic studies of this behavior. It possesses a rich vocal repertoire^5^ and exhibits turn-taking dynamics that closely parallel those observed in humans,^6^ making it a powerful primate model for probing the neural substrates of conversational coordination.

Turn-taking unfolds at least two separable temporal dimensions, sequence and rhythm. Sequence organizes information across successive acoustic elements, supporting the accumulation and integration of content. In humans, such integration may extend over timescales of minutes, with meaning gradually emerging as linguistic hierarchy unfold.^7–9^ Rhythm organizes the timing of exchange, enabling interlocutors to maintain interactional flow—for example, by adjusting response latency based on a partner’s intra-call and inter-call intervals.^3,10^ Together, these two dimensions imply that turn-taking is inherently multi-timescale, requiring the nervous system to link rapidly varying acoustic cues to longer-timescale contextual and interactive structure.

Current research in marmosets has not yet fully addressed how such multi-timescale integration is implemented, particularly under naturalistic, free-moving conditions. Existing marmosets research has largely emphasized behavior^10,11^ or neural activity confined to a limited set of brain regions.^12,13^ When large-scale neural measurements are available, analyses of acoustic feature processing are often performed under head-fixed conditions.^14^ This is a notable limitation: neuronal responses can differ between head-fixed and free-moving states,^15^ and marmosets typically vocalize substantially less during head fixation, constraining the very behavior of interest. Moreover, processing vocal information is unlikely to be implemented by a single locus, instead they are expected to depend on distributed, interacting brain regions.^14^ Therefore a critical question remains unresolved: how do marmosets integrate acoustic information across timescales to enable turn-taking, and how do distributed networks jointly encode sequential and rhythmic features in free-moving animals?

Answering this question requires large-scale functional measurements in fully free-moving animals. Conventional methods such as fMRI provide broad spatial coverage but severely constrain natural behavior. Electrophysiology can allow recordings in free-moving animals,^16^ yet typically samples a relatively limited spatial extent and often requires invasive implantation. Optical approaches can also be performed in free-moving animals,^17^ but remain restricted in field of view and depth. Taken together, these constraints highlight a practical need for a modality that can combine broad spatial coverage with use in unrestrained animals. Functional ultrasound imaging (fUS) offers a compelling alternative by combining high-coverage functional readout with practical compatibility for free-moving paradigms.

fUS provides high-coverage hemodynamic imaging and has been applied across species,^18–22^ but free-moving marmosets pose two challenges. First, it requires a durable, low-maintenance imaging interface that can deliver high-quality images while mitigating motion-related artifacts in unconstrained behavior. Second, it is necessary to reliably localize brain regions within images acquired in non-standard imaging planes, and to establish a robust preprocessing pipeline to suppresses locomotion-related contamination of the functional signal.

Here, we address these challenges by developing an exogenous marker– based fUS platform and a dedicated data-preprocessing pipeline. Using this platform, we show that intra-call prosodic perturbations of sequence and rhythm of vocal turn-takings elicit robust, distributed changes of neural responses. Collectively, our work establishes a practical platform for brain-wide functional imaging in free-moving animals and provides candidate brain regions and acoustic features that may contribute to marmoset conversational coordination, offering a foundation for future mechanistic studies.

## Results

### Development of functional ultrasound imaging pipeline in free-moving marmoset

Naturalistic, free-moving conditions impose two primary constraints on acquisition of large-scale ultrasonic brain signals: long-term cranial-window stability and resilience to locomotion-induced image degradation. An experimental platform was therefore developed to permit unrestricted behavior while preserving stable neurophysiological recording (Figure 1A).

**Figure 1.**
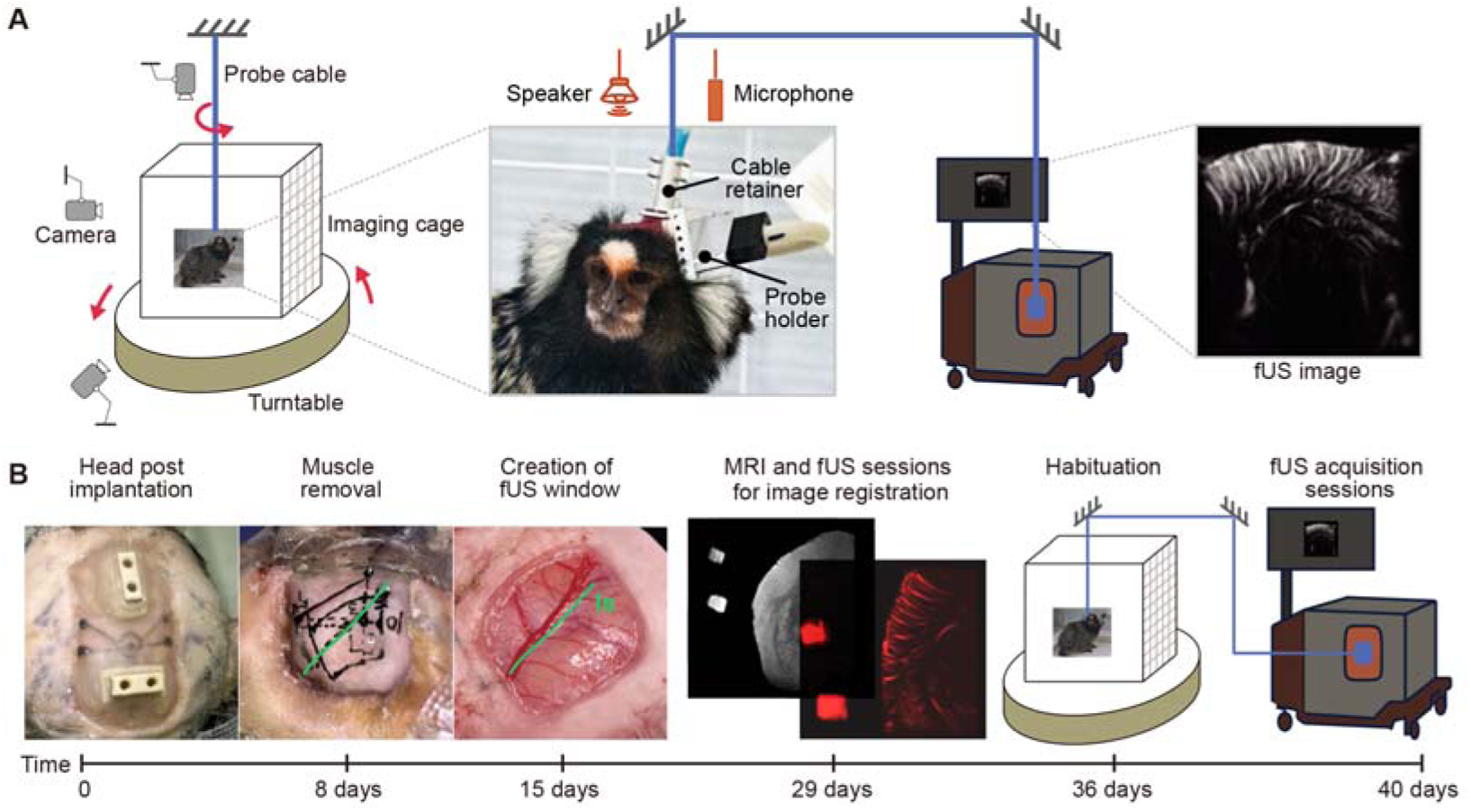
Overview of system setup and experimental workflow for functional ultrasound imaging in free-moving marmoset. (A) Overview of fUS setup for free-moving marmoset, including an imaging cage on an anti-twist turntable, three behavioral monitoring cameras, microphone and speaker, functional ultrasound imaging system. (B) Overview of experimental workflow, including head post implantation, muscle removal, creation of the fUS window, MRI and fUS sessions for image registration, habituation, and fUS acquisition sessions.

First, longitudinal recordings in free-moving marmosets require a durable cranial preparation (Figure 1B). A reliable cranial fUS window was established over the temporal lobe with accessories (Figure S1). This preparation remained functional for at least three months and up to more than one year and exhibited a low incidence of infection, enabling stable long-term measurements (Table 1 in supplementary information).

Second, rapid locomotion can introduce probe cable traction that compromises image quality. Cable tension may shift the TPX membrane relative to the surrounding skull, generating air cavities and producing signal dropout beneath the cavities (Figure S1F, red arrow). To mitigate this motion-related effect, an auxiliary head-mounted component (“Cable retainer” in Figure 1A) was designed to anchor the cable at the top of the head. This configuration reduced direct traction on the imaging window and eliminated the associated signal loss (Figure S1F, yellow arrow).

### Exogenous marker–based registration and optimized preprocessing to remove locomotion-related effects

Our experimental platform enabled stable, high-quality recordings in free-moving marmosets. However, brain regions cannot be reliably identified directly from non-standard imaging planes. To address this limitation, an exogenous marker-based, MRI-compatible registration workflow was established to register ultrasound images into template space (Figure 2; Figure S2).

**Figure 2.**
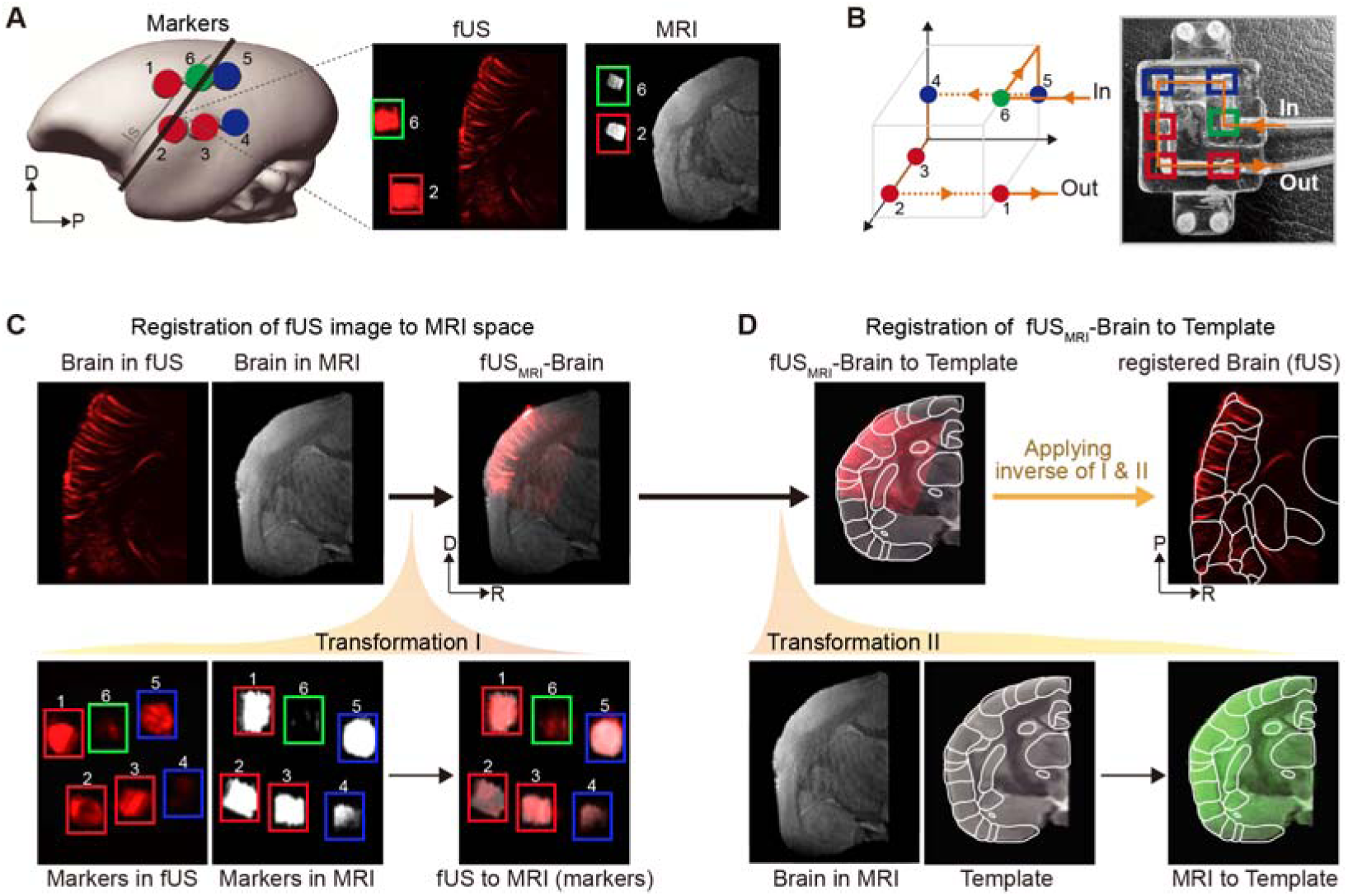
fUS image registration workflow through exogenous markers. (A) Exogenous markers visible in both MRI and fUS. Left panel: red, blue, and green dots indicate six markers, a bold black line indicates the fUS plane parallel to the lateral sulcus. Right panel: fUS image and MRI image showing brain slice and two of markers. (B) Design of flow-based exogenous marker system. Left panel: colored dots indicate six markers, with each color representing one depth. Yellow arrows indicate directions of fluid flow. Right panel: corresponding physical component. (C-D) Workflow of fUS image registration. (C) Transformation I is obtained by registering markers in fUS image to the same markers in MRI (bottom panel), and is applied to register fUS image to MRI (upper panel). (D) Transformation II is obtained by registering MRI image to template (bottom panel), and is applied to register fUS_MRI_ image to template (upper left). Inverse transformations are applied to project the template back onto fUS image (upper right).

The marker defines a 3D coordinate system using six fiducial points distributed across three mutually orthogonal planes at different depths (Figures 2A-B), enclosing the imaged brain within the coordinate system. In addition, a sealing component was introduced (Figure S2B) to prevent coupling-agent leakage and to improve imaging quality. To ensure cross-modality visibility, modality-specific fluids were used to delineate the tube’s inner contours (water for MRI and milk for fUS; Figure S2D).

Using the shared markers, transformation I was generated to register brain in ultrasound image to MRI space (fUS_MRI_-Brain, Figure 2C). Using the MRI brain image, transformation II was generated to register fUS_MRI_-Brain to template (Figure 2D). Using the inverses of transformations I and II, template-defined brain-region labels were projected back onto ultrasound space (Figure 2D, yellow arrow), yielding atlas-based regional annotations directly on ultrasound images. Following alignment, the markers and brain boundaries exhibited strong cross-modal correspondence, supporting accurate structural delineation.

Free-moving recordings introduced substantial locomotion-related contamination into ultrasound image signal: the animals’ locomotion not only directly perturbed the measurements, but also drove coupling-medium motion and induced relative displacement between the probe and the brain, thereby introducing locomotion-related fluctuations into the time series, encompassing both neural and non-neural components. To address this challenge, we implemented a preprocessing pipeline for the ultrasound data (Figure 3). The workflow comprised logarithmic transformation, spike regression, realignment, smoothing, skull-stripping, nuisance signal regression, and baseline drift removal (Figure 3A). Collectively, these steps substantially enhanced overall image quality (Figure 3B-C) and robustly attenuated locomotion-related contamination in the time series (Figure 3E).

**Figure 3.**
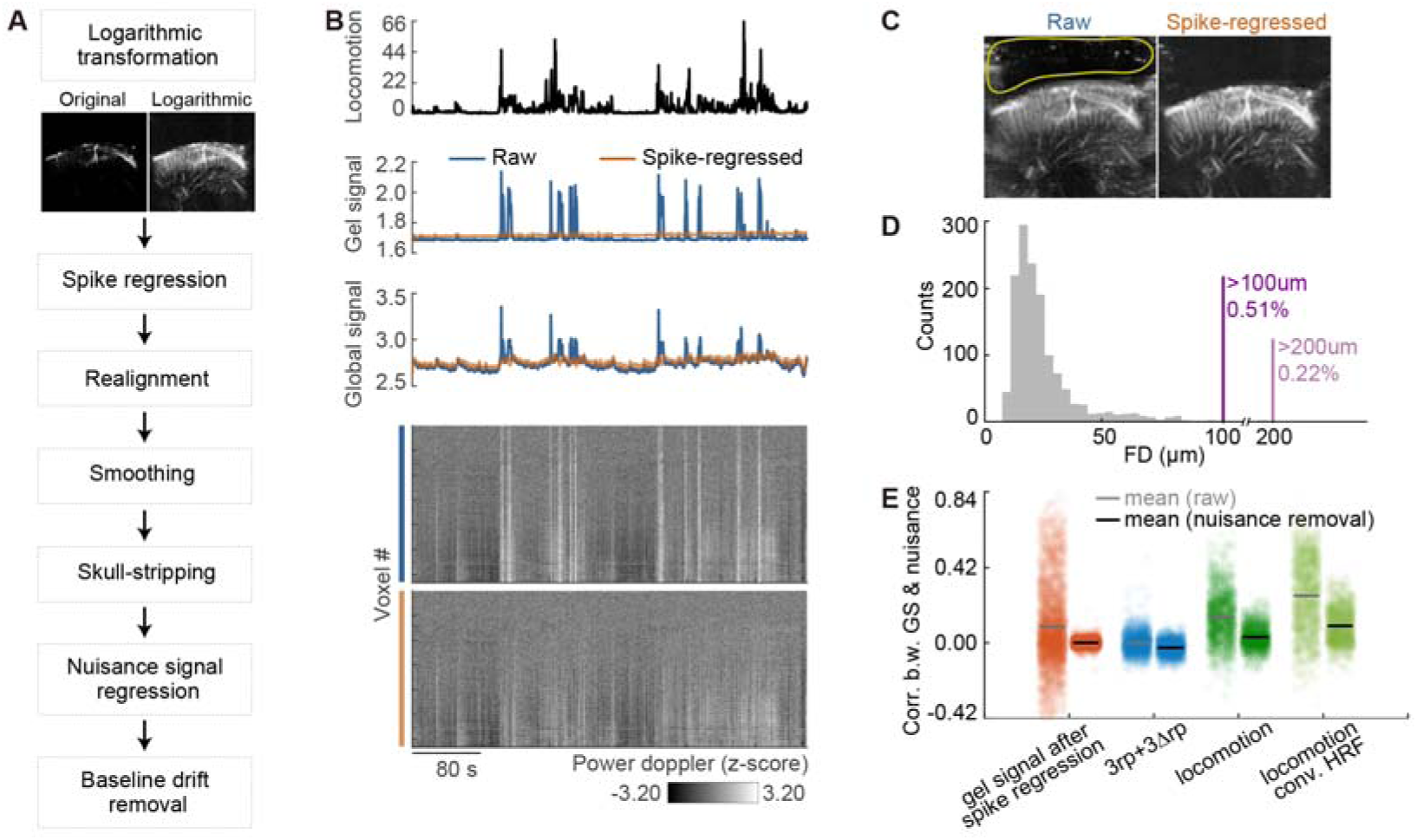
Functional ultrasound image preprocessing workflow. (A) Overview of the preprocessing pipeline, including logarithmic transformation, spike regression, realignment, smoothing, skull-stripping, nuisance signal regression, and baseline drift removal. (B) Spike regression suppresses high-intensity artifacts associated with large animal locomotion. From top to bottom: locomotion time course from an example run, raw signal (blue) from the coupling-gel region (yellow circle in (C)) and its spike-regressed version (orange), global brain signal (blue) and its spike-regressed version (orange), and a voxel-wise comparison of raw versus spike-regressed signals from the same run. (C) Example images at 104 s from the run shown in (B), displayed for the raw (left panel) and spike-regressed (right panel) data. (D) Distribution of frame-wise displacement (FD) between the fUS probe and the marmoset brain during acquisition. (E) Nuisance regressors including spike-regressed coupling-gel signal (orange), probe-brain displacement (3rp+3△rp, blue), animal locomotion (green), and locomotion convolved with a marmoset hemodynamic response function^37^ (light green).

### Functional validation of fUS pipeline using auditory tonotopy

To functionally validate the fUS workflow under free-moving conditions, we leveraged auditory tonotopy—a well-established organizing principle of the marmoset auditory system—as a validation paradigm. Functional ultrasound experiments were performed in marmosets B2 and B8 using 32 pure-tone stimuli spanning 440∼27,720 Hz (Figure 4A). In both animals, GLM analysis identified significant tone-evoked responses in auditory cortex (Figure 4B; Figure S3A; *p* < 0.05, FDR corrected).

**Figure 4.**
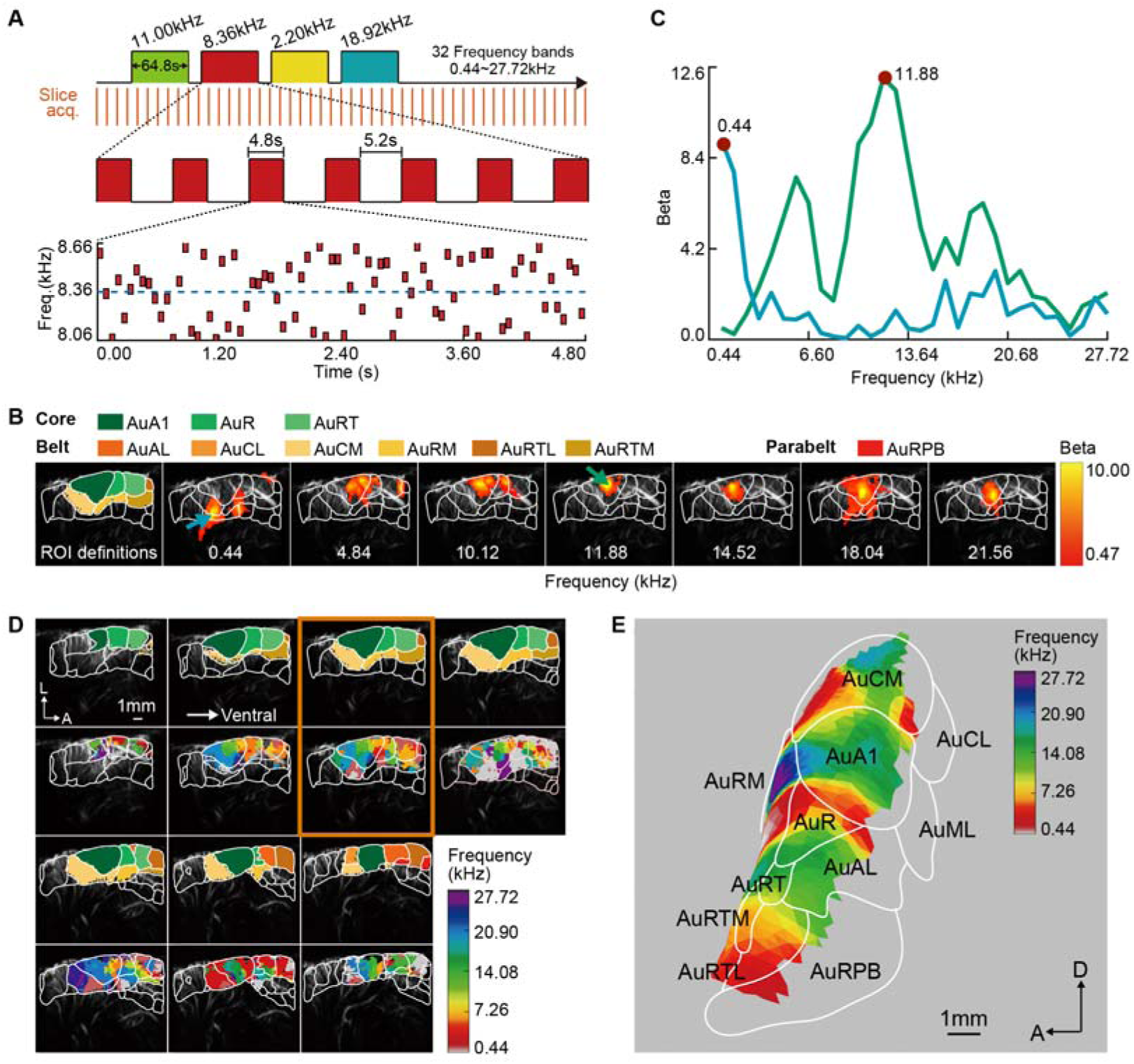
Validation of experimental pipeline through fUS-based tonotopy mapping in free-moving marmoset B2. (A) Stimulus design for tonotopy mapping using 32 frequency bands ranging from 440 to 27,720 Hz. Each frequency band is repeated 21 times. (B) Representative fUS response to frequency bands in a single slice (FDR corrected, *p*<0.05). White lines indicate boundaries of brain regions. Abbreviations of brain regions are listed in Table 2 in supplementary information. (C) Selective responses to low and middle frequency bands in regions indicated by blue and green arrows in (B). (D) Top panel: ROI definitions across all imaged slices after registration of template to fUS images. Bottom panel: tonotopy of each imaging slice. The colormap denotes the most responsive frequency (best-frequency) of each voxel. (E) Tonotopy in the auditory cortex after projecting volumetric results in (D) to the cortical surface.

Beyond overall responsiveness, multiple auditory subregions exhibited frequency-selective tuning, with systematic differences in best frequency across space (Figure 4C). Pixel-wise best-frequency maps were then computed for each imaging plane (Figure 4D; Figure S3B) and projected onto the cortical surface to reconstruct the tonotopy of auditory cortex (Figure 4E; Figure S3C). The resulting tonotopy recapitulated three hallmark features reported previously:^23^ (i) frequency reversals at established areal boundaries (AuRT-AuR and AuR-AuA1 in B2; AuRT, auditory cortex rostrotemporal, AuR, auditory cortex rostral area; AuA1, auditory cortex primary area); (ii) the expected anterior-low to posterior-high best-frequency gradient in AuA1; and (iii) a low-to-high frequency gradient extending from the anterior core toward the lateral belt of auditory cortex.

Together, these results support the reliability of the platform, including data acquisition in free-moving animals, atlas-based regional annotation, and preprocessing steps that effectively reduced locomotion-related confounds without disrupting canonical functional organization.

### Auditory and para-insular cortex selectively tracks intra-phrase sequence order

Interactive phee bouts provide a naturalistic substrate for studying how the temporal lobe decodes sequence and rhythm. To probe the relevant temporal integration window, a compact stimulus set was constructed to selectively perturb ordering and timing at multiple timescales. These stimuli were presented while functional ultrasound signals were recorded from the temporal lobe in three free-moving marmosets (Figure 5A).

**Figure 5.**
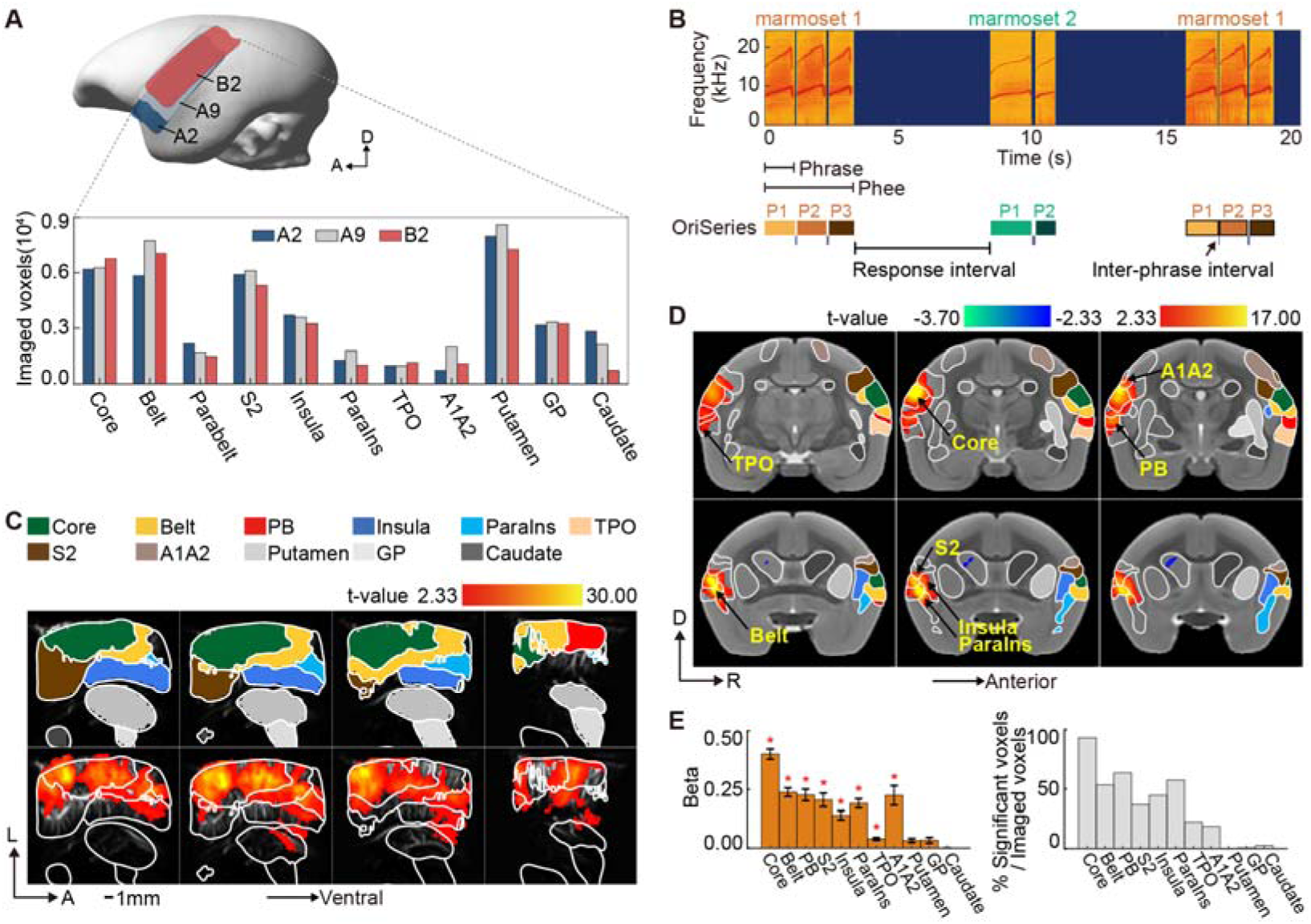
Original phee bouts induce robust activation in auditory-related regions across subjects. (A) Coverage of imaged regions for three marmosets (A2 in blue, A9 in gray, B2 in red), along with the number of imaged voxels per brain region for each animal. Abbreviations of brain regions are listed in Table 2 in supplementary information. (B) Auditory stimulation using original phee bouts (OriSeries). Each phee call comprises multiple phrases (P1, P2, P3) separated by inter-phrase intervals, and successive phee calls within a bout are separated by response intervals. (C) Top panel: ROI definitions. Bottom panel: Individual-level activation map elicited by OriSeries (*p* < 0.01, uncorrected), shown for subject A2. (D) Group-level activation map elicited by OriSeries (*p* < 0.01, uncorrected; *n* = 3). (E) Mean beta values (left panel, mean ± s.e.m) and significant voxel counts (right panel) (thresholded at *p* < 0.01, uncorrected). * Regions pass the group threshold and show a consistent response direction in all three marmosets.

After preprocessing, robust stimulus-evoked responses were observed across key auditory and associative structures (Figure S4; *p* < 0.05, FDR corrected). Original phee bouts (OriSeries, Figure 5B) elicited significant, distributed activation at the individual level, encompassing auditory core, belt, and parabelt regions (PB), insular and para-insular cortex (ParaIns), secondary somatosensory cortex (S2), areas 1 and 2 of cortex (A1/A2), and the temporo-parieto-occipital association region (TPO) (Figure 5C; Figure S5; *p* < 0.01, uncorrected). A highly similar spatial pattern was also evident at the group level (Figure 5D-E; *p* < 0.01, uncorrected, *n* = 3). Together, these results indicate that original phee bouts drive reproducible, multi-regional activation in free-moving animals, providing a foundation for decoding analyses of sequential and rhythmic manipulations.

Sensitivity to sequential perturbations at different timescales of the original phee bout was then assessed (Figure 6A). Among the three sequence manipulations, disruption of the internal ordering of phrase-level elements (SwapHalf) produced the most robust and reliable effect. Relative to OriSeries, SwapHalf elicited pronounced response enhancements in auditory core and belt regions as well as insular and para-insular cortex, evident both at the individual level (Figure S6; *p* < 0.01, uncorrected) and at the group level (Figure 6B-C; *p* < 0.01, uncorrected, *n* = 3). In contrast, SwapPhrase (reordering phrases within a call) and SwapPhee (reordering calls within the bout) generated strong responses in individual animals (Figures S7-8; *p* < 0.01, uncorrected), but yielded inconsistent and spatially restricted effects at the group level (Figure 6B-C; *p* < 0.01, uncorrected, *n* = 3). This dissociation suggests that phrase-level sequential structure provides the most consistent driver of population-level sensitivity in this dataset, whereas the other manipulations at longer timescale exhibit substantial inter-individual variability.

**Figure 6.**
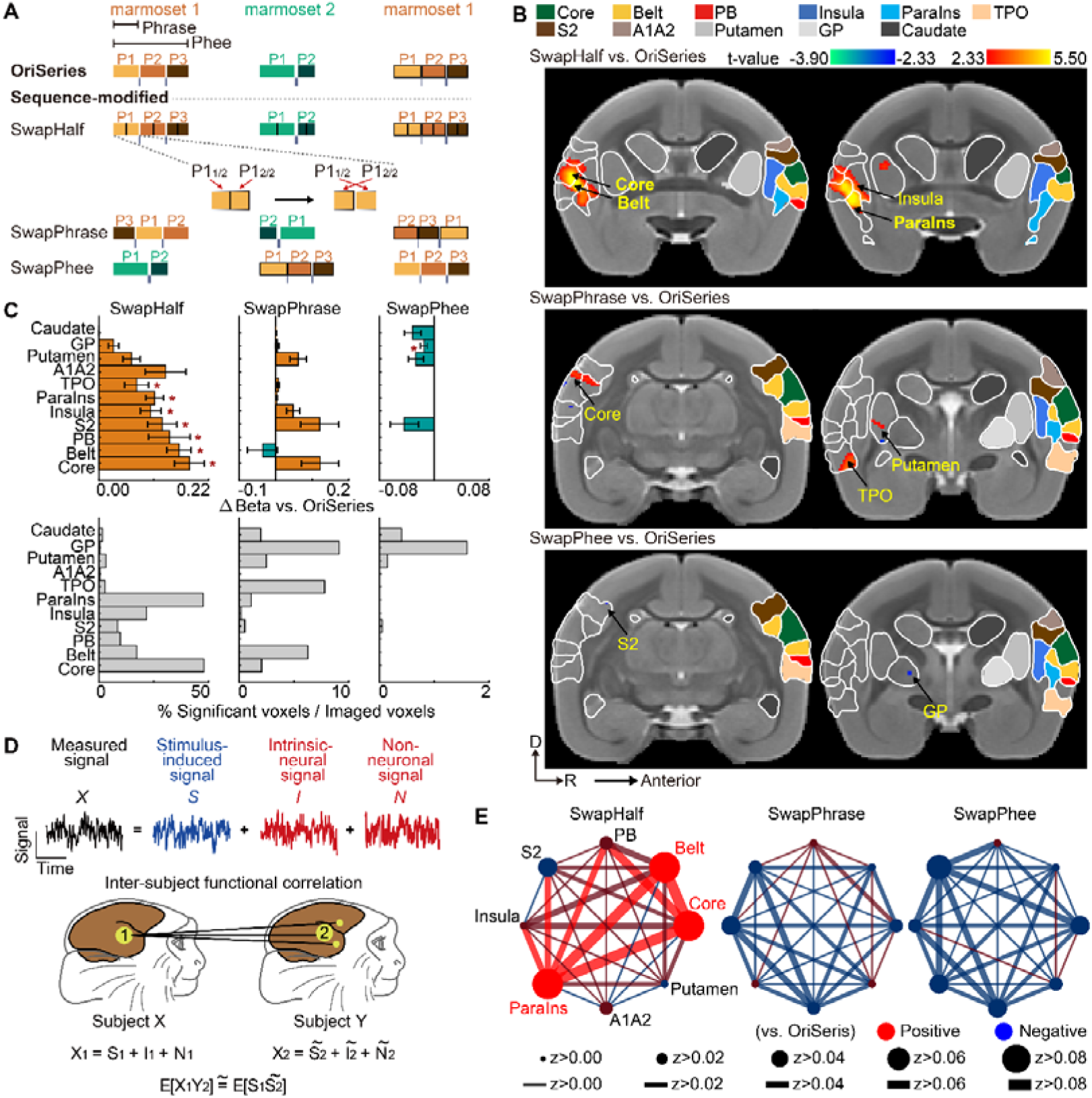
Auditory and para-insular regions are sensitive to intra-phrase sequence reordering but not to reordering at longer timescales. (A) Auditory stimulation design for three sequence-modified versions of the OriSeries across multiple timescales. SwapHalf: the first and second halves of each phrase are exchanged. SwapPhrase: the order of phrases within each phee is shuffled. SwapPhee: the order of phee calls within the vocal bouts is shuffled. (B) From top to bottom: Group-level contrast maps for SwapHalf vs. OriSeries, SwapPhrase vs. OriSeries, and SwapPhee vs. OriSeries (*p* < 0.01, uncorrected, *n* = 3). (C) Top panel: mean beta values for each modified version relative to OriSeries across regions (thresholded at *p* < 0.01, uncorrected), displayed as mean ± s.e.m. Bottom panel: significant voxel counts. * Regions pass the group threshold and show a consistent response direction in all three marmosets. (D) Schematic illustration of the principle of inter-subject functional correlation (ISFC) computation.^7^ (E) ISFC patterns for three sequence-modified versions relative to OriSeries. Filled circle indicates node strength; line width indicates pairwise correlation strength. Dark red (dark blue) nodes and edges represent enhanced (weakened) ISFC relative to the OriSeries. Red (blue) nodes and edges denote statistically significant increases (decreases) (*p* < 0.05, FDR corrected, *n* = 3).

To quantify cross-regional functional correlation sensitive to sequential manipulations, inter-subject functional correlation (ISFC) was employed, an approach that can isolate stimulus-locked dynamics shared across individuals (Figure 6D).^7^ For each animal, regional seeds were defined as those exceeding a response threshold (*p* < 0.05, uncorrected), followed by estimation of inter-regional ISFC. Similarly, SwapHalf significantly increased regional functional connection strength within auditory core, belt, and para-insular regions. Inter-regional connections, including core-belt, core-parabelt, core-para-insular, and belt-para-insular connections, were also enhanced (Figure 6E; *p* < 0.05, FDR corrected, *n* = 3). By comparison, SwapPhrase and SwapPhee did not yield ISFC changes that aligned with the GLM pattern, further supporting the conclusion that intra-phrase reordering evokes the most robust and cross-subject-consistent neural responses under free-moving conditions.

### Activity in the core region is sensitive to rhythmic changes at the inter-phrase timescale but not to longer timescale

Along the rhythmic dimension (Figure 7A), a directional, timescale-selective sensitivity to interval structure emerged. Shortening inter-phrase intervals (PhraseSpeedUp) increased responses in auditory core, whereas lengthening the intervals (PhraseSpeedDown) reduced responses (Figure 7B-C; Figures S9-10; *p* < 0.01, uncorrected). Beyond auditory core, PhraseSpeedUp also enhanced activity in para-insular cortex.

**Figure 7.**
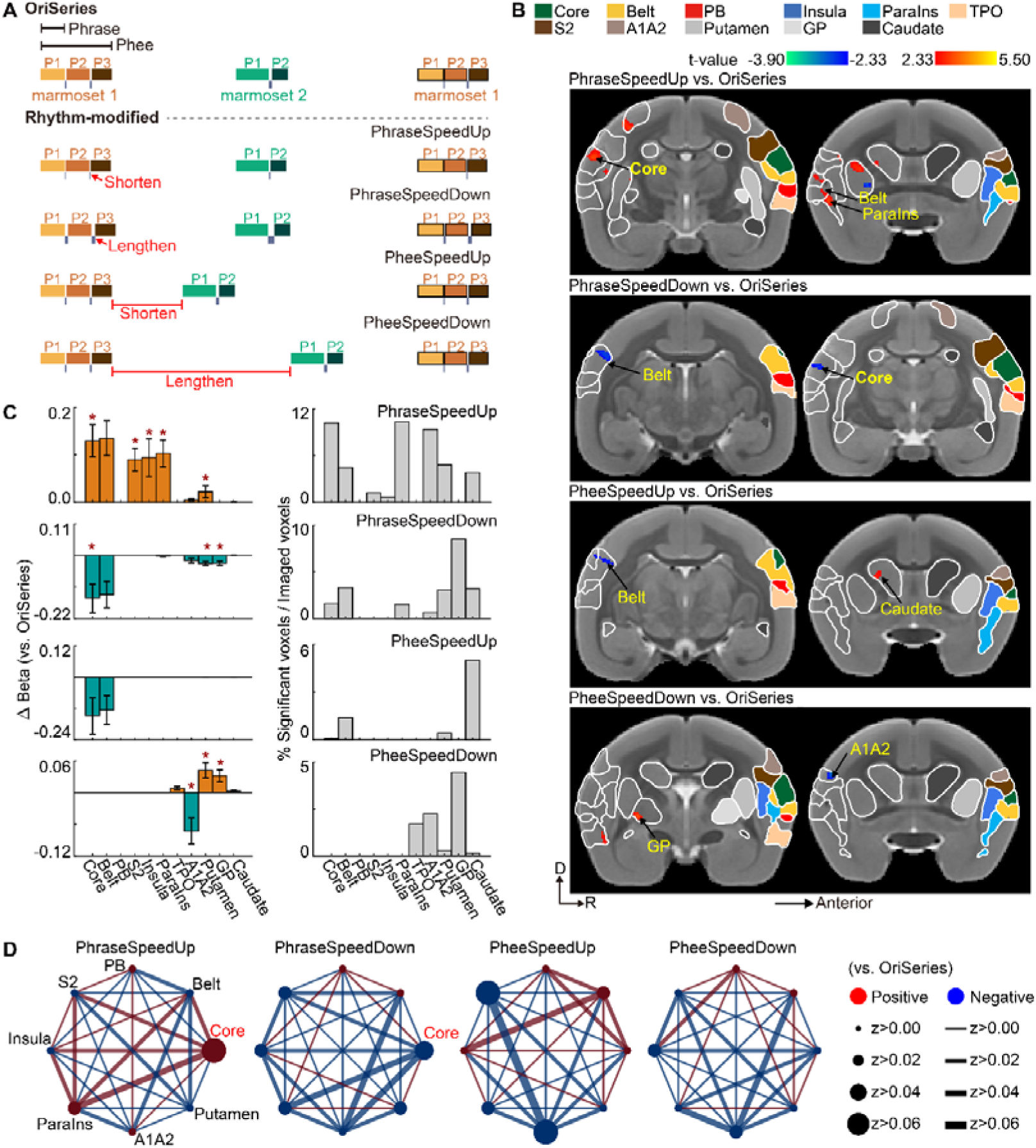
Activity in the core region is sensitive to rhythmic changes at the inter-phrase timescale but not to longer timescale. (A) Auditory stimulation design illustrating four rhythm-modified versions of the OriSeries across multiple timescales. The inter-phrase interval is shortened in PhraseSpeedUp and lengthened in PhraseSpeedDown. The response interval is shortened in PheeSpeedUp and lengthened in PheeSpeedDown. (B) From top to bottom: Group-level contrast maps for PhraseSpeedUp vs. OriSeries, PhraseSpeedDown vs. OriSeries, PheeSpeedUp vs. OriSeries, PheeSpeedDown vs. OriSeries (*p* < 0.01, uncorrected, *n* = 3). (C) Top panel: mean beta values for each modified version relative to OriSeries across regions (thresholded at *p* < 0.01, uncorrected), displayed as mean ± s.e.m. Bottom panel: significant voxel counts. (D) ISFC patterns for the four rhythm-modified versions relative to OriSeries.

Converging evidence from connectivity supported this rhythmic effect. ISFC analysis revealed interval-locked coupling changes centered on auditory core, consistent with modulation of auditory-core dynamics by inter-phrase interval perturbations (Figure 7D; *p* < 0.05, FDR corrected, *n* = 3).

By comparison, although modest effects were detected in some individuals for shortened response interval (PheeSpeedUp; Figure S11; *p* < 0.01, uncorrected) and lengthened response interval (PheeSpeedDown; Figure S12; *p* < 0.01, uncorrected) these effects were spatially weak at the group level and showed limited concordance between the univariate (GLM) and ISFC readouts (Figure 7C-D). Collectively, these findings indicate that neural responses during vocal interaction are reliably tuned to rhythmic changes at the phrase-level timescale, but not to longer timescale.

## Discussion

In this study, we extended functional ultrasound imaging to unrestrained marmosets and established a platform including external marker-based registration workflow and data-preprocessing pipeline. Using this platform, we found that when marmosets listened to vocal bouts, they were especially sensitive to intra-call perturbations of sequence and rhythm.

Since its introduction to neuroscience,^24^ fUS has been adopted across a wide range of species.^18–22^ Naturally, it is necessary to conduct regional delineation for functional ultrasound images. Prior work has relied on vascular signatures,^18^ imaging planes anchored to anatomical landmarks (e.g., sulci/gyri),^25^ the intrinsic functional architecture of specific areas,^22^ or clustering voxels according to the similarity of their temporal activity profiles.^21^ Despite their utility in well-controlled acquisition regimes, reliable anatomical labeling remains a major barrier in primate fUS when recordings are constrained to local, non-standard planes, where landmark visibility can be limited. In this study, we addressed this limitation through an exogenous marker-based, bimodal registration workflow that achieved accurate fUS-MRI structural alignment (Figure 2). Importantly, this approach is broadly applicable: it infers spatial transformation information for arbitrary imaging planes without relying on specific anatomical landmarks or functional references, and is therefore especially suited to localized imaging in non-standard planes.

Free-moving fUS has been demonstrated in rodents,^19,26^ but marmosets’ rapid, ballistic movements lead to more severe locomotion artifacts. We modeled and corrected key motion-related errors, including coupling-medium fluctuations and probe–brain relative motion (Figure 3). As a result, our preprocessing framework is designed to remain effective in imaging paradigms characterized by frequent and vigorous movements, extending the applicability of free-moving fUS beyond low-motion conditions. Collectively, these hardware and analytical advances increased the robustness and physiological interpretability of free-moving hemodynamic imaging in marmosets and were validated in the auditory tonotopy experiment (Figure 4). Using this platform, we address a long-standing question in auditory neuroscience: over what temporal windows do animals extract sequential and rhythmic structure from natural vocal communication? By combining naturalistic stimulation with broad spatial coverage, our results dissociate contributions of intra-call sequence organization and interaction timing to auditory cortical responses.

Phrase-internal reordering elicited the most consistent responses across subjects (Figure 6), suggesting a phrase-level integration window for parsing natural sequences. Compared with prior studies with limited coverage,^12,13^ responses extended beyond core auditory cortex into belt and para-insular regions, indicating a distributed network for sequence processing. These effects may reflect sensitivity to relative pitch dynamics, as phrase-level reordering necessarily alters pitch progression across the phrase, consistent with evidence that superior temporal cortex encodes relative pitch variation.^27^ However, the direction of our effect differs from reports of reduced spiking following phrase-level reordering, a discrepancy that may reflect differences in the regions examined,^12^ call types tested,^13^ and measurement modality (fUS versus spiking), which can diverge in sign and spatial integration due to vascular and population-level contributions.

Along the rhythmic dimension, our findings indicate that auditory core responses track the tempo of conversational exchange: shorter inter-phrase intervals were associated with enhanced responses, whereas longer intervals attenuated activity (Figure 7). This dependence on interval duration supports the view that rhythmic timing is dynamically regulated during social interaction rather than being a fixed property of an individual’s vocal output. Convergent interaction-dependent timing adjustments have been reported across taxa, including antiphonal singing birds (slowed syllable production accompanied by reduced HVC firing)^3^ and singing mice (delayed syllable onsets and lengthened syllables upon partner entry),^1^ and behavioral work in marmosets suggesting shorter inter-phrase intervals during interaction than in isolation.^28^ Collectively, these convergent observations support the possibility that interaction-dependent regulation of vocal rhythm is a conserved strategy, with early auditory circuits tracking these temporal contingencies to facilitate turn-taking and coordination.

More broadly, the sequence- and rhythm-related signatures identified here map onto key building blocks of prosody, which integrates pitch dynamics, syllable duration, inter-phrase intervals, and related temporal cues to support prediction in communicative exchange. Developmental findings further underscore the predictive value of prosodic cues for turn-taking,^29,30^ particularly when lexical information is insufficient.^29^ Outside humans, evidence for semantic information in vocalizations is confined to a small number of taxa, whereas vocal signals in most species carry little clearly referential content,^31,32^ therefore prosodic structure may represent a conserved substrate for anticipating turn boundaries and coordinating vocal timing. In this context, our combination of naturalistic paradigms and expanded spatial coverage provides a framework for localizing and quantifying how auditory systems extract intra-call sequential and rhythmic structure in real-world communication.

This study had several limitations that warrant further investigation. First, recordings were confined to the temporal lobe. Frontal lobe also contributes to vocal processing^14^ and supports integration over longer timescales.^33^ Stimuli that yielded low response consistency here may elicit more stable responses in frontal lobe, which were not assessed. Second, subjects acted as third-party listeners rather than engaged interactants, potentially reducing attention to the auditory input. Third, respiration and heart rate were not recorded, precluding assessments of arousal state. In free-moving contexts, some marmosets did not tolerate thoracic restraint (e.g., belt-mounted wireless devices) or added weight, and such restraint markedly suppress vocalization. Therefore, miniaturized recording device is much needed in the future.

Taking together, these advances establish an extensible platform for future studies of dyadic marmoset interactions. This approach can be readily integrated with diverse neuromodulation methods to delineate the circuit mechanisms supporting neural function under naturalistic conditions. Using this platform, we further demonstrate that intra-call manipulations of sequence and rhythm robustly modulate neural responses, highlighting these features—and their underlying substrates—as promising targets for mechanistic investigation.

## Methods

### Animals

Data were obtained from four adult male common marmosets (age 3∼5 years; body weight 350∼420 g). Animals B2 (name: Xiaomei), A2 (Xiaonengmiao), and A9 (Wuding) participated in experiment “Probing timescales of sequence and rhythm in phee bouts”. Animals B2 and B8 (Tiaotiaotang) participated in the experiment “Auditory tonotopy”. The marmosets were housed in the primate facility under controlled temperature (27∼30°C) and a 12 h:12 h light-dark cycle. During experimental sessions, animals had access to ample water and received small food rewards. All procedures were approved by the Animal Care and Use Committee at the Center for Excellence in Brain Science and Intelligence Technology, Chinese Academy of Sciences (Ethics approval number: CEBSIT-2020023).

### Surgical procedures

All surgical procedures were performed under standard aseptic conditions. Following a 12-hour fast, marmosets received preoperative subcutaneous injections 1 hour before surgery: atropine (to reduce glandular secretions, 0.15 mg/kg), dexamethasone (to prevent cerebral edema, 0.5 mg/kg), etamsylate (to reduce bleeding, 46.88 mg/kg), and meloxicam (to prevent infection, 10 mg/kg). Throughout surgery, heart rate, respiration, body temperature, oxygen saturation, perfusion pressure, anesthetic concentration, and oxygen flow were continuously monitored to ensure stable anesthesia. To minimize surgical burden and improve tolerance, the whole procedure was conducted in three following steps, spaced one week apart.

***Head post implantation***: Under isoflurane anesthesia, the marmoset was placed prone in a stereotaxic frame. A midline sagittal incision was made along the scalp to expose the skull, and the periosteum was removed. 6∼8 PEEK screws (M1.4 × 0.6 mm) were inserted around the planned head-post site. The head post (Figure 1B, Figure S1A) was positioned approximately 3 mm posterior to the bregma and reinforced with dental acrylic (LANG, USA). Postoperatively, meloxicam oral suspension (0.1 mg/kg) was administered orally once daily for three consecutive days.

***Temporal muscle removal***: Preoperative medications were identical to those used for head post implantation, with the addition of intramuscular flunixin meglumine (25 mg/kg). The animal was positioned laterally with the left temporal muscle facing upward. Local anesthesia with lidocaine was applied to the surgical site. The temporal muscle was bluntly dissected from the skull and then rapidly transected using an electrocautery pen to expose the temporal bone, providing swift hemostasis and a clear operative field. After thorough hemostasis and irrigation of residual muscle tissue with saline, the muscle stump was sealed with medical adhesive (Beijing Fuaile Technology Development Co., Ltd.), and a thin protective layer of dental acrylic was applied to the skull (Figure 1B). Postoperatively, flunixin meglumine and meloxicam oral suspension were administered for three consecutive days.

***Cranial window creation for ultrasound imaging***: Preoperative medications were identical to those used for head post implantation. After the imaging window area was delineated, the skull within the defined region was removed. Because a major vessel typically runs within the meninges near the lateral fissure in this region, and preserving it often resulted in postoperative hemorrhage, a partial meningeal resection was performed. The dura and arachnoid were carefully separated from the pia using forceps and a nasal dissector; the dura and arachnoid were then removed, preserving only the pia mater. A TPX film (thickness 0.125 mm; Goodfellow, UK) was trimmed to fit and bonded tightly to the surrounding skull with 401 adhesive (Loctite, Germany) to seal the window. During film application, a small amount of saline was dripped onto the pia to maintain tissue moisture. The primary headpiece (Figure S1B) was fixed to the skull around the window with dental acrylic; after the acrylic cured, a transparent protective cap (Figure S1C) was secured to the primary headpiece. Postoperatively, meloxicam oral suspension and dexamethasone tablets (0.23 mg/kg) were administered for three consecutive days.

### Imaging apparatus for free-moving marmosets

***Imaging cage***: The imaging cage, serving as the marmoset’s activity space, was constructed from five joined panels of transparent acrylic (polymethyl methacrylate, PMMA). Iron climbing grids were installed on the right and rear walls. The bottom was not enclosed with acrylic, instead, a rotatable turntable was installed. The turntable was connected to a remote controller via an infrared cable. When the ultrasound probe cable on the marmoset’s head became excessively twisted, the turntable was activated to rotate slowly in the reverse direction, thereby relieving the entanglement. The internal dimensions of the cage were 32 × 32 × 38 cm^3^.

***Behavior recording***: Three cameras (Zhongweiaoke, model A1080P) were positioned at the front, left, and top of the imaging cage to record the marmoset’s behavior, at distances of 31, 29, and 18 cm from the cage, respectively. Video was recorded at 30 fps with a frame size of 1280 × 720 pixels. Three LEDs were installed around the cage, controlled by an Arduino. LEDs was switched on at the beginning of each experimental run and switched off at the end. We analyzed animal’s behavior only during the illuminated period.

***Stimuli delivering***: A speaker (MF1, TDT) was placed approximately 90 cm from the cage to deliver auditory stimuli.

***Vocalization recording***: A microphone (378C01, PCB, TDT; frequency range 4∼100,000 Hz) was mounted on the top of the cage to record marmoset vocalizations during imaging.

### Functional ultrasound imaging system

***fUS parameters***: The fUS platform (Iconeus, France) utilized a 15 MHz linear probe with 128 elements (pitch: 0.11 mm; bandwidth: 60%). The focal depth was 8 mm. Spatial resolution was 0.10 × 0.11 mm^2^ (axial × lateral). The head-mounted flexible cable measured 43 cm and weighed 21.4 g. Structural imaging parameters: image matrix 128 × 183 × 193, voxel size 0.11 × 0.10 × 0.10 mm^3^, field of view (FOV) 14 × 20 mm^2^ (width × depth), frame rate 2.5 Hz. Functional imaging parameters: image matrix 128 × 193, voxel size 0.11 × 0.10 mm^2^, slice thickness 0.4 mm. Structural imaging was acquired to provide reference data for image registration. Functional imaging was acquired for the experiment “Auditory tonotopy” and the experiment “Probing timescales of sequence and rhythm in phee bouts”.

The imaged volume was centered on the auditory cortex. Data were acquired as a multi-slice series with the scan orientation aligned to the course of the lateral sulcus. Each cranial window encompassed multiple imaging planes. Depending on window size and the experimental paradigm, 6∼22 slices were defined per marmoset, spaced at 0.3∼0.5 mm intervals. Only a single slice was acquired per imaging session, the complete slice set for a given animal was assembled across multiple days.

***Probe fixation***: The ultrasound probe was securely mounted above the targeted brain region of the free-moving marmoset using a custom-designed fixation assembly. The assembly comprised five components (Figure S1): a head post, a primary headpiece, a base, a probe holder F/M, and a cable retainer. The head post was fabricated using polyether ether ketone (PEEK), and all other components were 3D-printed using photosensitive resin. The total mass of the assembled device was 9.6 g.

The head post and primary headpiece were surgically affixed to the skull at the cranial vertex and around the cranial window, respectively. During imaging, the base was secured to the primary headpiece with screws. The probe was then clamped between the F and M halves of the probe holder and mounted onto the base. Different imaging planes were obtained by adjusting the distance between the holder’s cutting plane and a predefined reference point. Finally, the cable retainer was attached to the head post, and the flexible segment of the probe cable was secured to the retainer.

### Registering ultrasound image to marmoset template

***Data acquisition for cross-modal image registration***: Two weeks after surgery, each marmoset underwent one MRI session and one fUS session. The registration device consisted of two components: a core module and an auxiliary module. The core module contained an internal channel with six markers (Figure 2C; Figure S2A, colored dots), which served as shared exogenous fiducials visible in both fUS and MRI for image registration. The auxiliary module (Figure S2B) was designed to prevent leakage of the coupling medium during scanning, thereby improving imaging quality.

MRI experiments were conducted in a 9.4T scanner (Bruker, Germany). After induction of anesthesia with isoflurane, the core module was mounted onto the color-coded holes of the primary headpiece (Figure S2C). The internal channel of the core module was filled with water, and the inlet and outlet were clamped after complete filling. A single-loop RF coil (20-mm diameter; Bruker) was attached to the exterior of the core module for MRI acquisition. MR images were acquired using FcFLASH sequence with the following parameters: matrix 300 × 350 × 250, field of view 30 × 35 × 25 mm³, isotropic resolution 0.1 × 0.1 × 0.1 mm³, echo time (TE) 8.73 ms, repetition time (TR) 50 ms, flip angle 7°. The resulting MR images included both the marmoset brain and the six markers (hereafter MRI-Brain and MRI-Markers, respectively).

After MRI acquisition, the RF coil was removed, and the auxiliary module (Figure S2B) was mounted onto the primary headpiece at the same color-coded positions (Figure S2C) to prevent loss of the acoustic coupling medium during fUS acquisition. The water in the device channels was drained and replaced with milk solution (20 g milk powder in 180 mL water), which was circulated at a constant flow rate using a pump. Using the structural ultrasound settings described in the “Functional ultrasound imaging system” section, we first acquired ultrasound images of the six markers (fUS-Markers). The core module was then removed while the animal brain remained in the same position, and ultrasound images of the marmoset brain were acquired using identical parameters (fUS-Brain).

***Registration of ultrasound images to the standard marmoset template***: Using ITK-SNAP (https://www.itksnap.org), we generated binary masks for the six markers from MRI-Markers and fUS-Markers, and masks for the marmoset brain from MRI-Brain and fUS-Brain (Figure 2A). All subsequent processing was performed on data extracted using these masks. We first registered fUS-Markers to MRI-Markers using ANTs antsRegistration (https://www.nitrc.org/projects/ants), yielding Transformation I. Transformation I was then applied with antsApplyTransforms to align fUS-Brain to MRI-Brain (fUS_MRI_-Brain). Next, MRI-Brain was registered to a standard marmoset template^34^ to obtain Transformation II. Applying Transformation II to align the fUS_MRI_-Brain to template. Transformation I included an affine matrix, whereas Transformation II comprised an affine matrix and a nonlinear warp field. Using the inverse of Transformations I and II, the standard template was also warped back into fUS-Brain space.

### Experimental design

***Auditory tonotopy*:** Using MATLAB R2018b (MathWorks, USA), we generated 32 frequency bands spanning 440∼27,720 Hz with uniform spacing. Adjacent band center frequencies were separated by 880 Hz. Stimuli were generated at a 96-kHz sampling rate. Each band consisted of 80 pure tones centered on the band center frequency with a jitter of ± 300 Hz. Each tone lasted 0.05 s and was separated by an inter-tone interval of 0.01 s, resulting in a total duration of 4.80 s per band. The inter-band interval was 5.20 s. Each band was presented 21 times, with band order randomized. For animal B2, seven imaging planes were acquired (total recording time, 800.80 min). For animal B8, six imaging planes were acquired (total recording time, 686.40 min).

***Probing timescales of sequence and rhythm in phee bouts:*** Auditory stimuli were obtained from three breeding pairs of common marmosets housed at the marmoset facility of the Center for Excellence in Brain Science and Intelligence Technology, Chinese Academy of Sciences. We selected 33 phee bouts (termed OriSeries) recorded during natural vocal exchanges in which pair members were visually occluded from one another. Phee calls, which consist of multiple phrases, are commonly used for long-distance communication. Each bout contained 4∼6 phee calls, and the response latency between partners ranged from 1 to 12 s.

We generated three sequence-manipulated versions relative to OriSeries by altering temporal structure at three timescales. (1) Intra-phrase timescale, termed SwapHalf: each phrase was split into two halves, and the two halves were exchanged. (2) Intra-call timescale, termed SwapPhrase: within each phee call, the order of phrases was randomly permuted while the position of each phee call within the bout was preserved. (3) Inter-call timescale, termed SwapPhee: the internal phrase structure of each phee call was preserved, but the order of phee calls within the bout was permuted.

We also generated four rhythm-manipulated versions relative to OriSeries at the intra-call and inter-call timescales. At the intra-call timescale, inter-phrase intervals within each phee call were either reduced by 90% or increased twofold, yielding PhraseSpeedUp and PhraseSpeedDown, respectively. At the inter-call timescale, the inter-call intervals produced by the first marmoset in OriSeries were kept unchanged, whereas the response latency of the second marmoset to the first was decreased or increased by 50%, yielding PheeSpeedUp and PheeSpeedDown, respectively.

Within each run, OriSeries and each of the seven modified versions were presented once. Each stimulus lasted 42 s, separated by an 8∼12 s inter-stimulus interval. Condition order was randomized within each run. The total duration of each run was 460 s.

Animals A2, A9, and B2 contributed 14, 18, and 22 imaging planes, respectively. For each imaging plane, 33 runs (corresponding to the 33 phee bouts) were acquired. Total recording times were 3,542 min (A2), 4,554 min (A9), and 5,566 min (B2). Because some imaging planes were obtained from only one or two animals, all analyses were restricted to 14 planes from A2, 15 from A9, and 13 from B2 except Figure S4.

### Preprocessing of functional ultrasound images

The preprocessing pipeline included logarithmic transformation, spike regression, realignment, smoothing, and skull stripping (Figure 3A). For analyses requiring ISFC, preprocessing additionally included nuisance signal regression and baseline drift removal. Preprocessing was implemented using custom scripts in MATLAB R2018b and SPM12 (https://www.fil.ion.ucl.ac.uk/spm).

***Spike regression***: To suppress high-amplitude spike artifacts (Figure 3B), we delineated the coupling-gel region in the images (Figure 3C, yellow circle) using a mask derived from ITK-SNAP. We extracted the gel-region time series and performed principal component analysis (PCA). The first ten principal components were used as reference signals and regressed out from the voxel-wise image time series, thereby removing spike-like transients coupled to the gel signal.

***Realignment***: Motion correction was performed using SPM12 (Realign). Because the data were acquired as a single slice, only three nonzero realignment parameters were estimated (translation in two in-plane directions and in-plane rotation; referred to as 3 rp). Framewise displacement (FD) was computed for each scan assuming a marmoset brain radius of 15 mm.^35^ Runs with a maximum FD exceeding 100 µm were excluded (Figure 3D).

***Smoothing***: Spatial smoothing was applied using an anisotropic Gaussian kernel to match the anisotropic acquisition resolution (FWHM: 0.20 mm axial, 0.22 mm lateral, and 0.80 mm elevational).

***Nuisance signal regression (ISFC preprocessing only)***: To reduce task-unrelated variance (Figure 3E), four classes of nuisance signals were regressed out.

***(1). Slow gel-related fluctuations***. Gradual shifts in the coupling gel were modeled using the top two principal components (PC2) from a PCA of the coupling-gel region.

***(2). Marmoset behavior***. We calibrated a three-camera setup using MATLAB R2018b (stereoCameraCalibrator) to recover each camera’s pose parameters. We then used DeepLabCut v2.3.9 (https://www.mackenziemathislab.org/deeplabcut) to extract behavioral time series from videos recorded from three viewpoints. Because of frequent occlusions, only the head was labeled, which yielded the highest tracking accuracy. The 2D estimates were combined with the calibration parameters using pose3D^36^ to reconstruct 3D head coordinates over time.

Because the behavioral tracking and fUS acquisition had different sampling rates, we computed the total head displacement within the corresponding time window for each fUS frame, yielding a locomotion-magnitude time series matched to the fUS time series. This series was convolved with the marmoset hemodynamic response function (HRF)^37^ to generate an HRF-modulated locomotion regressor. Both the raw locomotion-magnitude series and its HRF-convolved counterpart were included as nuisance covariates.

***(3). Probe-brain relative displacement***. Artifacts due to relative displacement between the ultrasound probe and brain were modeled using the three realignment parameters (3 rp) and their first temporal derivatives (3 Δrp).

***(4). Spontaneous vocalizations during imaging***. Marmosets occasionally vocalized during imaging. Vocalization onsets and offsets were identified using a dual-threshold endpoint detection method that combines short-time energy and the short-time average zero-crossing rate. We then constructed a binary time series (1 during vocalization; 0 otherwise) aligned to the imaging time course, and convolved it with the marmoset HRF.^37^

***Baseline drift removal (ISFC preprocessing only)***: To remove low-frequency drift, we applied a first-order Butterworth high-pass filter to the ultrasound time series (cutoff period: 128 s; sampling rate: 2.5 Hz).

### Data analysis

***Auditory tonotopy*:** We applied transformations I and II to warp the 32 GLM-derived activation maps from each of the two marmosets (one per frequency band) into the standard template space. For each voxel, we then defined its best frequency as the band showing the maximum activation magnitude, yielding a best frequency map in template space. Finally, we projected this volumetric map onto the cortical surface using the wb_command volume-to-surface-mapping tool (https://humanconnectome.org) with the -ribbon-constrained option.

***General linear model analysis*: *(1). Subject-level analysis***: Analyses were conducted in SPM12. Standard first-level analyses were performed on all single-slice runs. Of the four nuisance classes described in “Preprocessing of functional ultrasound images”, the first three were included as regressors in the design matrix, while spontaneous vocalizations and the experimental stimuli of interest were modeled as conditions. ***(2). Group-level analysis***: For each condition, beta images from the subject-level analyses were spatially registered to a marmoset template. To mitigate registration error, the registered beta maps were smoothed with an 0.6 mm isotropic Gaussian kernel. A linear mixed effect model was subsequently used to generate activation maps for each condition and contrast maps for between-condition comparisons.

***Inter-subject functional correlation analysis*:** ISFC was calculated following a previous study.^7^ Briefly, ISFC measures stimulus-locked connectivity by correlating regional time courses across different subjects listening the same stimulus (Figure 6D). Because intrinsic fluctuations and most noise are not shared across individuals, ISFC suppresses these components and emphasizes stimulus-driven coupling, improving SNR and yielding more specific, replicable patterns than standard within-subject functional connectivity. we defined regions of interest (ROIs) as brain regions that surpassed the statistical threshold across all conditions (*p* < 0.05, uncorrected). We then computed ISFC among all ROI pairs. Multiple comparisons across condition-wise contrasts were controlled using false discovery rate correction.

## Data availability

The drawings for probe fixation and image registration, as well as the datasets used for image registration, are available at https://doi.org/10.12412/BSDC.1770895419.30001, and will be released to public upon publication.

## Code availability

Codes for image registration and data preprocessing are available at https://doi.org/10.12412/BSDC.1770895419.30001.

All raw data and analyses supporting the current study are available from the corresponding author on request.

## Author contributions

X.C., L.W. and Z.L. designed and supervised the study; X.C., B.W. performed the marmoset surgeries; B.W., M.C., S.W. collected the fUS data; X.C., B.B., K.Z. collected the MRI data; M.P. optimized the MRI sequence; X.C., Y.L., L.W. and Z.L. carried out the analysis and organized the results; X.C. and Z.L. wrote the original draft and revised the draft.

## Acknowledgements

The authors thank Marmoset and Mouse Facility of Center for Excellence in Brain Science and Intelligence Technology (Institute of Neuroscience) for animal care, and Brain Science Data Center, Chinese Academy of Sciences for support with data analysis and dataset release. This work was supported by the Strategic Priority Research Program of Chinese Academy of Sciences (XDB0930202 to Z.L.), National Natural Science Foundation of China (Excellent Young Scientists Fund 32322036 to Z.L.).

## Reference

1. Okobi, D.E., Banerjee, A., Matheson, A.M.M., Phelps, S.M., and Long, M.A. (2019). Motor cortical control of vocal interaction in neotropical singing mice. Science 363, 983–988. 10.1126/science.aau9480.

2. Coleman, M., Day, N., Rivera-Parra, P., and Fortune, E. (2021). Neurophysiological coordination of duet singing. Proc. Natl. Acad. Sci. USA 118, e2018188118. 10.1073/pnas.2018188118.

3. Hoffmann, S., Trost, L., Voigt, C., Leitner, S., Lemazina, A., Sagunsky, H., Abels, M., Kollmansperger, S., Maat, A.T., and Gahr, M. (2019). Duets recorded in the wild reveal that interindividually coordinated motor control enables cooperative behavior. Nat. Commun. 10, 2577. 10.1038/s41467-019-10593-3.

4. Grijseels, D., Fairbank, D., and Miller, C. (2024). A model of marmoset monkey vocal turn-taking. Proc. R. Soc. B 291, 20240150. 10.1098/rspb.2024.0150.

5. Bezerra, B., and Souto, A. (2008). Structure and usage of the vocal repertoire of callithrix jacchus. Int. J. Primatol. 29, 671–701. 10.1007/s10764-008-9250-0.

6. Borjon, J., and Ghazanfar, A. (2014). Convergent evolution of vocal cooperation without convergent evolution of brain size. Brain Behav. Evol. 84, 93–102. 10.1159/000365346.

7. Simony, E., Honey, C., Chen, J., Lositsky, O., Yeshurun, Y., Wiesel, A., and Hasson, U. (2016). Dynamic reconfiguration of the default mode network during narrative comprehension. Nat. Commun. 7, 12141. 10.1038/ncomms12141.

8. Lyu, B., Choi, H., Clarke, A., Randall, B., and Tyler, L. (2019). Neural dynamics of semantic composition. Proc. Natl. Acad. Sci. USA 116, 21318-21327. 10.1073/pnas.1903402116.

9. Chang, C.H.C., Nastase, S., and Hasson, U. (2022). Information flow across the cortical timescale hierarchy during narrative construction. Proc. Natl. Acad. Sci. USA 119, e2209307119. 10.1073/pnas.2209307119.

10. Takahashi, D., Narayanan, D., and Ghazanfar, A. (2013). Coupled oscillator dynamics of vocal turn-taking in monkeys. Curr. Biol. 23, 2162–2168. 10.1016/j.cub.2013.09.005.

11. Pomberger, T., Risueño Segovia, C., Löschner, J., and Hage, S. (2018). Precise motor control enables rapid flexibility in vocal behavior of marmoset monkeys. Curr. Biol. 28, 788–794. 10.1016/j.cub.2018.01.070.

12. Jia, G., Bai, S., Lin, Y., Wang, X., Zhu, L., Lyu, C., Sun, G., An, K., Wang, A., Li, X., and Gao, L. (2023). Representation of conspecific vocalizations in amygdala of awake marmoset. Natl. Sci. Rev. 10, nwad194. 10.1093/nsr/nwad194.

13. Zeng, H., Huang, J., Li, J., Shen, Z., Gong, N., Wen, Y., Wang, L., and Poo, M. (2021). Distinct neuron populations for simple and compound calls in the primary auditory cortex of awake marmosets. Natl. Sci. Rev. 8, nwab126. 10.1093/nsr/nwab126.

14. Jafari, A., Dureux, A., Zanini, A., Menon, R., Gilbert, K., and Everling, S. (2023). A vocalization-processing network in marmosets. Cell Rep. 42, 112526. 10.1016/j.celrep.2023.112526.

15. Jovanovic, V., Fishbein, A., Mothe, L., Lee, K.-F., and Miller, C. (2022). Behavioral context affects social signal representations within single primate prefrontal cortex neurons. Neuron 110, 1318–1326.e1314. 10.1016/j.neuron.2022.01.020.

16. Tsunada, J., Wang, X., and Eliades, S. (2024). Multiple processes of vocal sensory-motor interaction in primate auditory cortex. Nat. Commun. 15, 3093. 10.1038/s41467-024-47510-2.

17. Zong, W., Obenhaus, H., Skytøen, E., Eneqvist, H., Jong, N., Jorge, M., Moser, M.-B., and Moser, E. (2021). Large-scale two-photon calcium imaging in freely moving mice. Cell 185, 1240–1256.e1230. 10.1016/j.cell.2022.02.017.

18. Grillet, M., Brunner, C., Macé, E., Montaldo, G., Urban, A., and roska, b. (2021). Whole-brain functional ultrasound imaging in awake head-fixed mice. Nat. Protoc. 16, 3547–3571. 10.1038/s41596-021-00548-8.

19. Urban, A., Dussaux, C., Martel, G., Brunner, C., Macé, E., and Montaldo, G. (2015). Real-time imaging of brain activity in freely moving rats using functional ultrasound. Nat. Methods 12, 873–878. 10.1038/nmeth.3482.

20. Dizeux, A., Gesnik, M., Ahnine, H., Blaize, K., Arcizet, F., Picaud, S., Sahel, J., Deffieux, T., and Pouget, P. (2019). Functional ultrasound imaging of the brain reveals propagation of task-related brain activity in behaving primates. Nat. Commun. 10, 1400. 10.1038/s41467-019-09349-w.

21. Zhang, Y.S., Takahashi, D.Y., El Hady, A., Liao, D.A., and Ghazanfar, A.A. (2022). Active neural coordination of motor behaviors with internal states. Proc. Natl. Acad. Sci. USA 119, e2201194119. 10.1073/pnas.2201194119.

22. Bimbard, C., Demené, C., Girard, C., Radtke-Schuller, S., Shamma, S., and Boubenec, Y. (2018). Multi-scale mapping along the auditory hierarchy using high-resolution functional UltraSound in the awake ferret. eLife 7, e35028. 10.1101/249417.

23. Song, X., Guo, Y., Li, H., Chen, C., Lee, J., Zhang, Y., Schmidt, Z., and Wang, X. (2022). Mesoscopic landscape of cortical functions revealed by through-skull wide-field optical imaging in marmoset monkeys. Nat. Commun. 13, 2238. 10.1038/s41467-022-29864-7.

24. Macé, E., Montaldo, G., Cohen, I., Baulac, M., Fink, M., and Tanter, M. (2011). Functional ultrasound imaging of the brain. Nat. Methods 8, 662–664. 10.1038/nmeth.1641.

25. Norman, S., Maresca, D., Christopoulos, V., Griggs, W., Demené, C., Shapiro, M., and Andersen, R. (2020). Single trial decoding of movement intentions using functional ultrasound neuroimaging. Neuron 109, 1554–1566.e1554. 10.1016/j.neuron.2021.03.003.

26. Sieu, L.-A., Bergel, A., Tiran, E., Deffieux, T., Pernot, M., Gennisson, J.-L., and Cohen, I. (2015). EEG and functional ultrasound imaging in mobile rats. Nat. Methods 12, 831–834. 10.1038/nmeth.3506.

27. Tang, C., Hamilton, L., and Chang, E. (2017). Intonational speech prosody encoding in the human auditory cortex. Science 357, 797–801. 10.1126/science.aam8577.

28. Gong, N., Huang, J., Ma, H., Sun, Y., and Chang, L. (2022). Complex rules of vocal sequencing in marmoset monkeys. Preprint at bioRxiv. 10.1101/2022.08.03.502601.

29. Casillas, M., and Frank, M.C. (2013). The development of predictive processes in children’s discourse understanding. Proc. Annu. Meet. Cogn. Sci. Soc. 35, 299–304. https://hdl.handle.net/11858/00-001M-0000-0013-F72F-6.

30. Shi, J., Gu, Y., and Vigliocco, G. (2022). Prosodic modulations in child - directed language and their impact on word learning. Dev. Sci. 26, e13357. 10.1111/desc.13357.

31. Owren, M., and Bernacki, R. (1988). The acoustic features of vervet monkey alarm calls. J. Acoust. Soc. Am. 83, 1927–1935. 10.1121/1.396477.

32. Slocombe, K., and Zuberbühler, K. (2005). Functionally referential communication in a chimpanzee. Curr. Biol. 15, 1779–1784. 10.1016/j.cub.2005.08.068.

33. Ding, N., Melloni, L., Zhang, H., Tian, X., and Poeppel, D. (2015). Cortical tracking of hierarchical linguistic structures in connected speech. Nat. Neurosci. 19, 158–164. 10.1038/nn.4186.

34. Liu, C., Yen, C., Szczupak, D., Tian, X., and Glen, D. (2021). Marmoset Brain Mapping V3: population multi-modal standard volumetric and surface-based templates. NeuroImage 226, 117620. 10.1016/j.neuroimage.2020.117620.

35. Power, J., Barnes, K., Snyder, A., Schlaggar, B., and Petersen, S. (2012). Spurious but systematic correlations in functional connectivity MRI networks arise from subject motion. NeuroImage 59, 2142–2154. 10.1016/j.neuroimage.2011.10.018.

36. Sheshadri, S., Dann, B., Hueser, T., and Scherberger, H. (2020). 3D reconstruction toolbox for behavior tracked with multiple cameras. J. Open Source Softw. 5, 1849. 10.21105/joss.01849.

37. Hirano, Y., Yen, C., Liu, J., Mackel, J., Merkle, H., Nascimento, G., and Stefanovic, B. (2017). Investigation of the BOLD and CBV fMRI responses to somatosensory stimulation in awake marmosets (Callithrix jacchus). NMR Biomed. 31, e3864. 10.1002/nbm.3864.

