## Supplemental Information for "Functional ultrasound imaging in free-moving marmosets reveals that intra-call prosodic features modulate large-scale cortical dynamics"

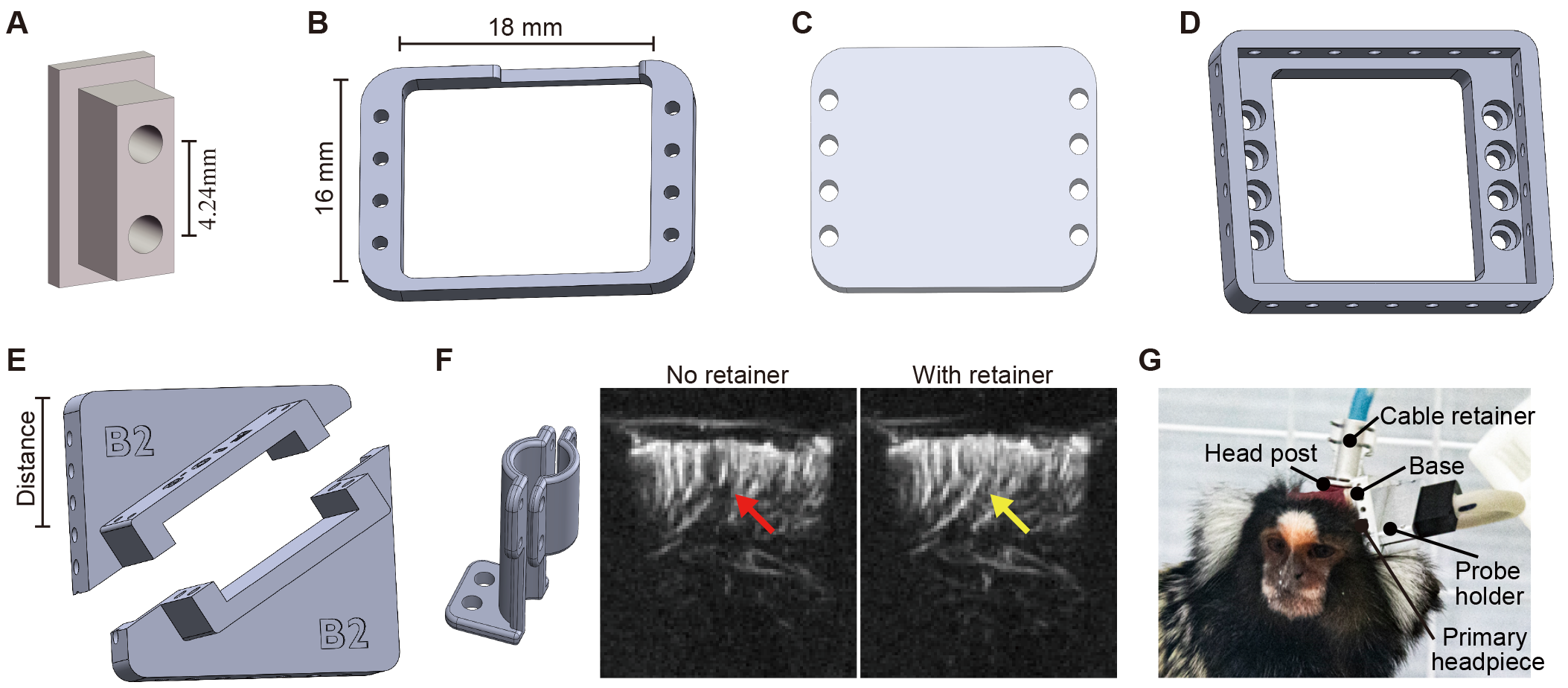

**Figure S1. Accessories for stabilizing and securing the ultrasound probe, related to Figure 1**

(A) Head post.

(B) Primary headpiece.

(C) Protective cap.

(D) Base.

(E) Probe holder F/M.

(F) Cable retainer (left) and improvement in image quality after installation (right). The red arrow indicates signal dropout, and the yellow arrow indicated signal recovery.

(G) Schematic of a marmoset wearing the complete set of fixation accessories.

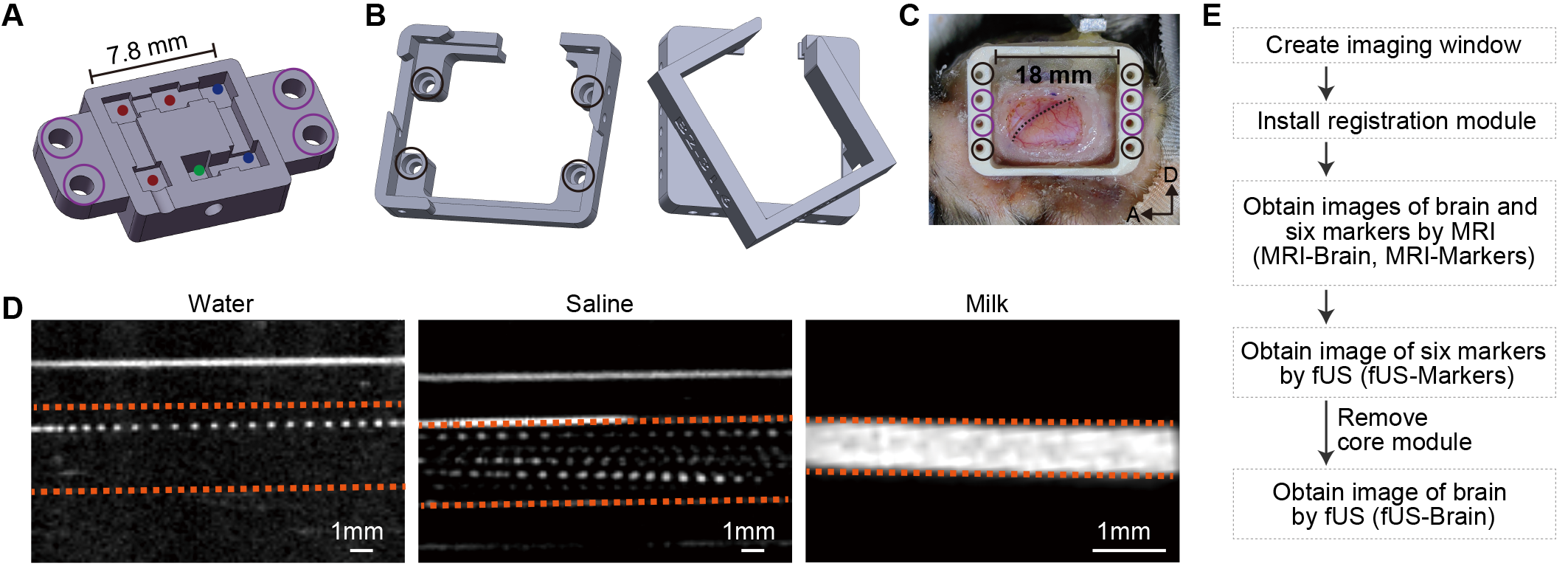

**Figure S2. Auxiliary components and fluid selection to improve fUS registration quality, related to Figure 2**

(A) Cross-sectional schematic of the core module showing internal channels. Colored dots (red, blue, green) denote markers at different depths. Purple circles indicate positions that lock into the correspondingly colored holes of the primary headpiece shown in panel (C).

(B) Auxiliary sealing module, designed to prevent leakage of the coupling gel and thereby maintain high-quality imaging. Black circles indicate positions that fix into the matching holes on the primary headpiece in panel (C).

(C) Imaging window with the primary headpiece installed. Black dashed line indicates the lateral sulcus.

(D) Comparison of image quality obtained with three types of fluid: water, saline, and milk. Orange dashed lines delineate the inner contour of the flow channel.

(E) Workflow for acquisition of registration data.

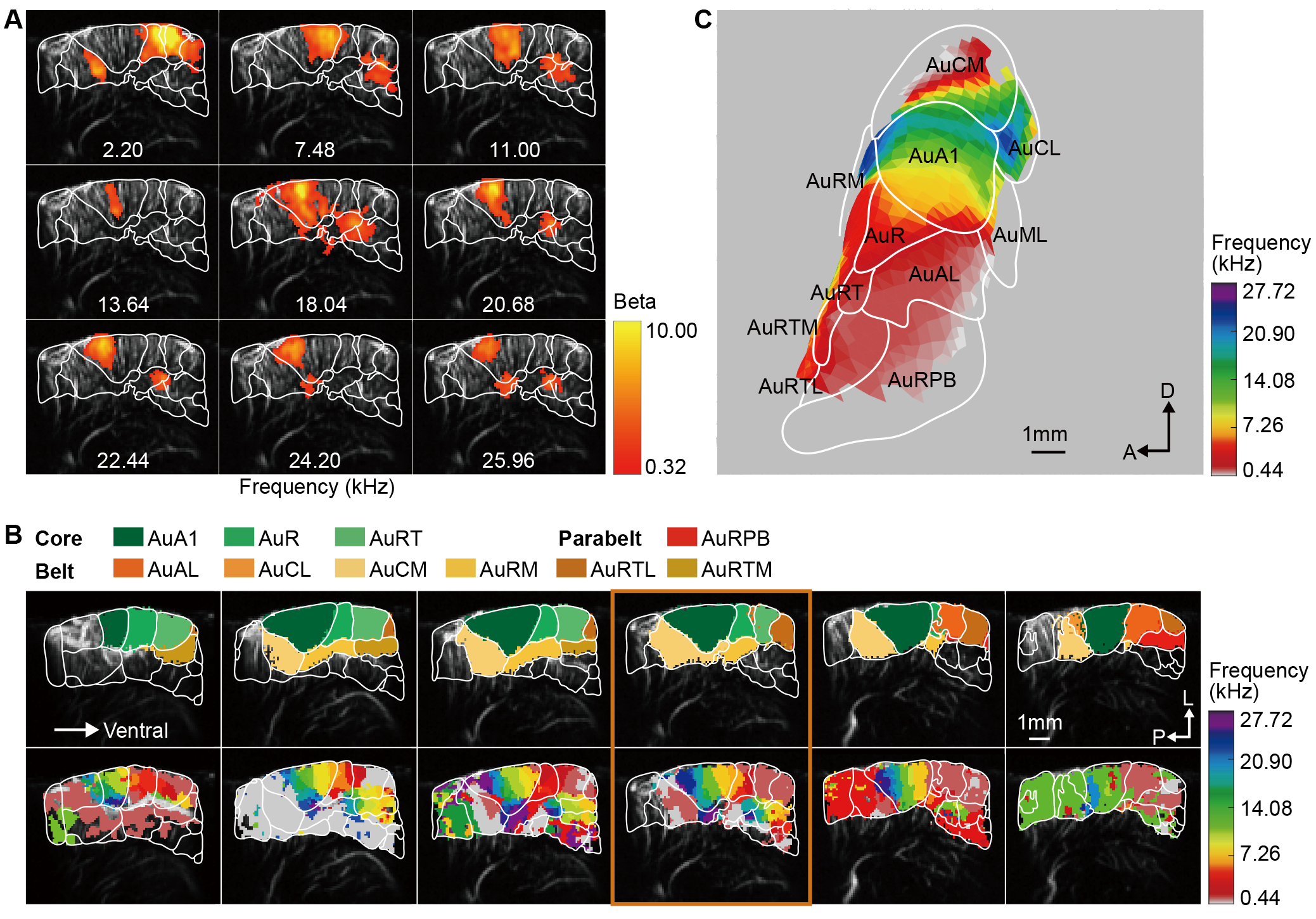

**Figure S3. Tonotopic organization in the auditory cortex of marmoset B8, related to Figure 4**

(A) Representative fUS response to frequency bands in a single slice (yellow box in panel (B), FDR corrected *p*<0.05). White lines indicate regional boundaries.

(B) Top panel: brain region definitions across all imaged slices after registration of template to fUS images. Bottom panel: tonotopy for each imaging slice. The colormap denotes the most responsive frequency of each voxel. Abbreviations of brain regions are listed in the Table 2 in supplementary information.

(C) Tonotopy in the auditory cortex after projecting volumetric results in (B) to the cortical surface.

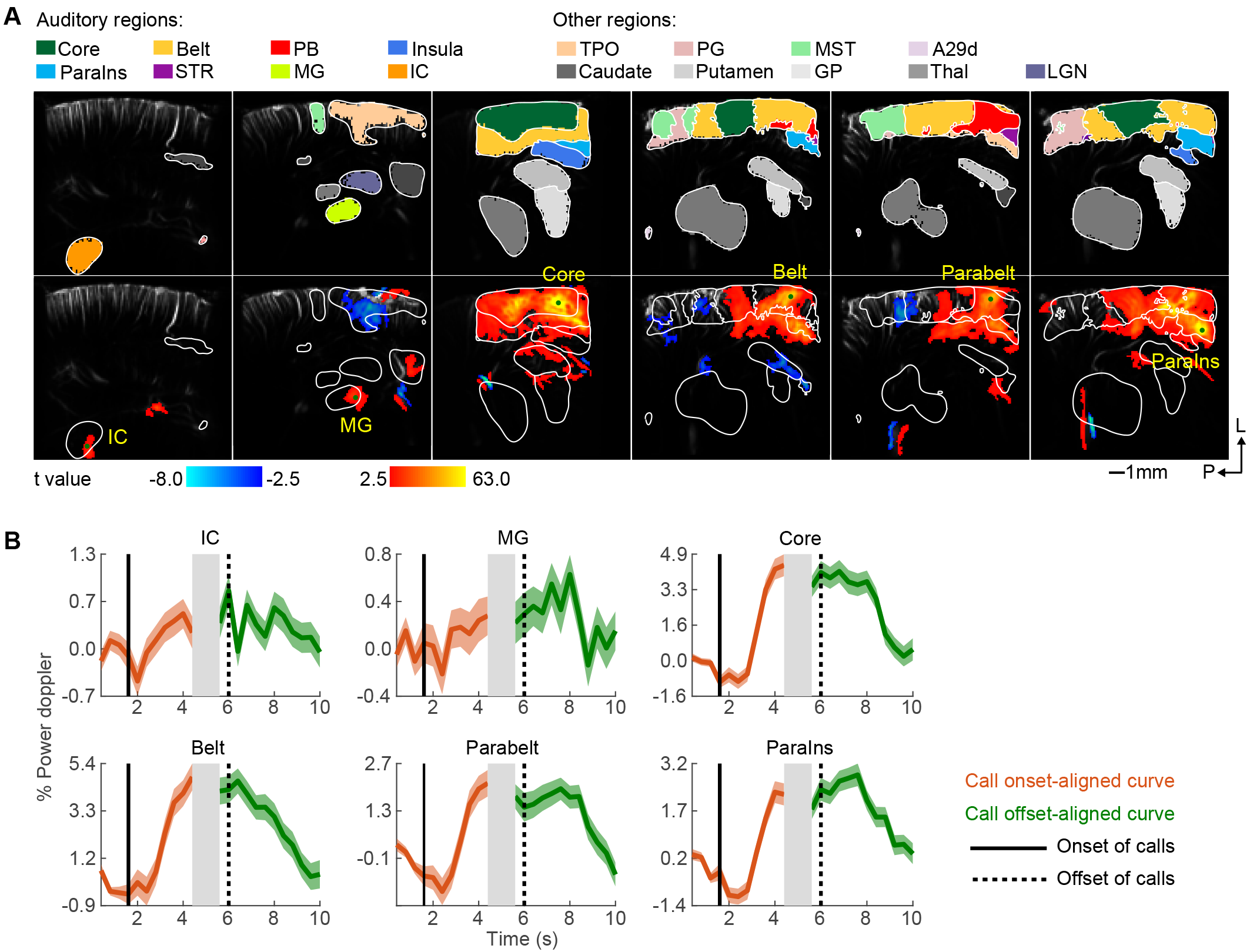

**Figure S4. Activation of the auditory pathway by phee calls, related to Figure 5**

(A) Top panel: brain region definitions. Bottom panel: auditory-related regions significantly activated by phee calls (GLM; *p* < 0.05, FDR corrected). Example activation map from marmoset B2.

(B) Event-aligned response time courses (orange and green) time-locked to onset (solid black line) and offset (dashed black line) of phee calls, shown as mean ± s.e.m across calls. Traces are extracted from the location marked by the green point in (A).

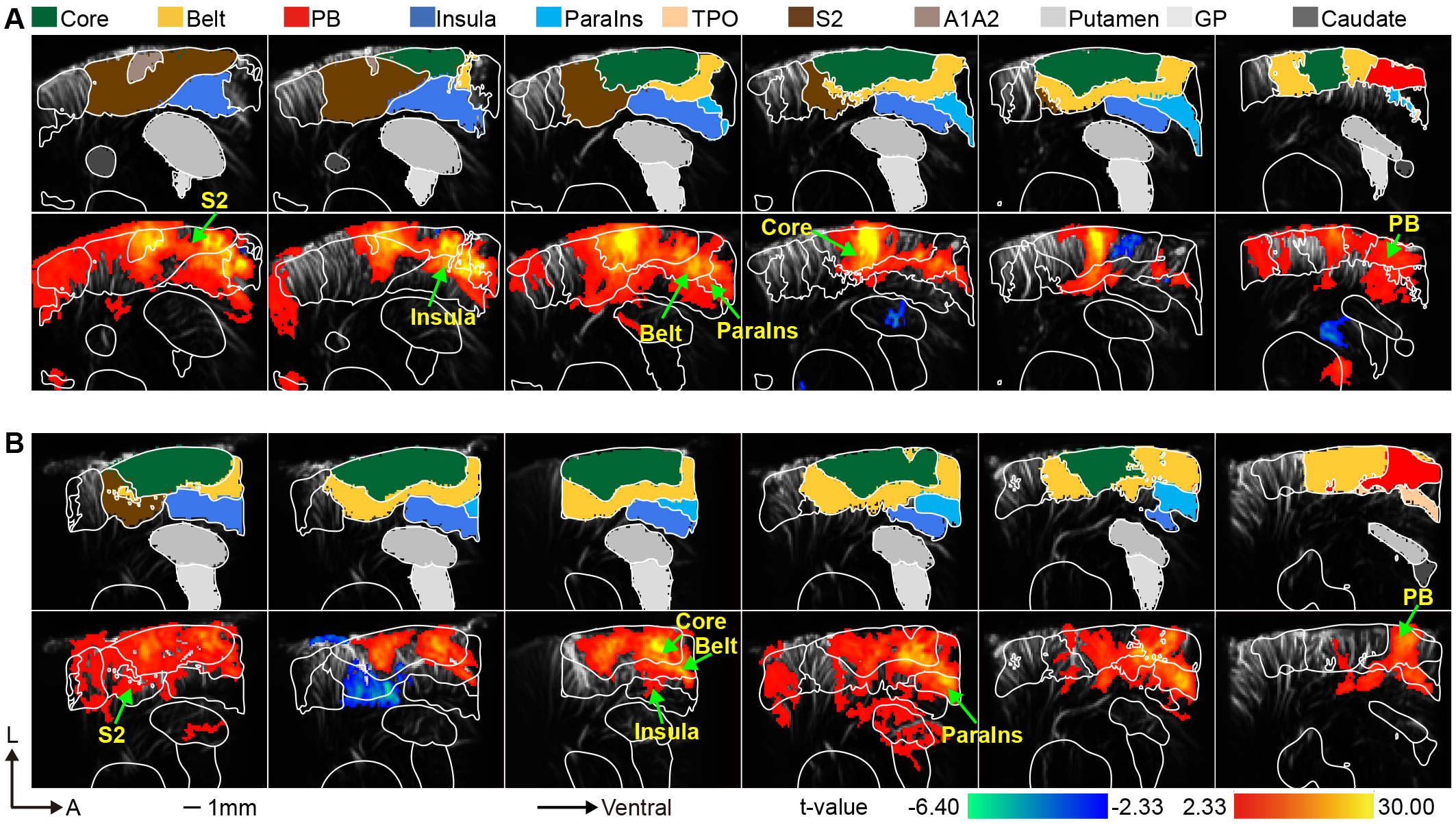

**Figure S5. Individual-level activation maps evoked by OriSeries, related to Figure 5**

(A) Top panel: brain region definitions. Bottom panel: activation map for marmoset A9 in response to OriSeries (GLM, *p* < 0.01, uncorrected).

(B) Activation map for marmoset B2 computed using the same analysis settings as in (A).

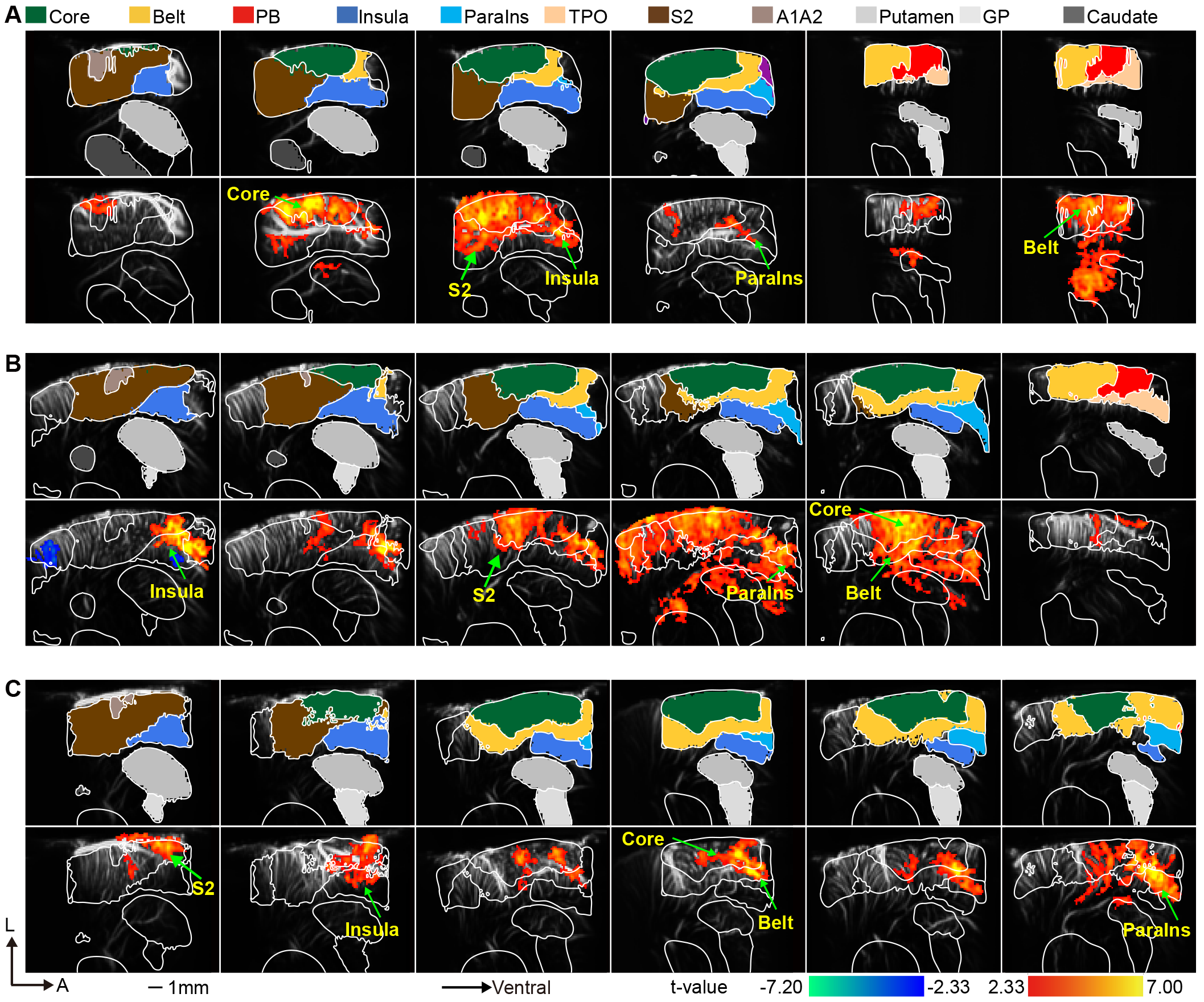

**Figure S6. Individual-level contrast maps evoked by SwapHalf vs. OriSeries, related to Figure 6**

(A) Top panel: brain region definitions. Bottom panel: contrast map (SwapHalf vs. OriSeries) for marmoset A2 (GLM, *p* < 0.01, uncorrected).

(B-C) Contrast maps for marmosets A9 (B) and B2 (C) computed using the same analysis settings as in (A).

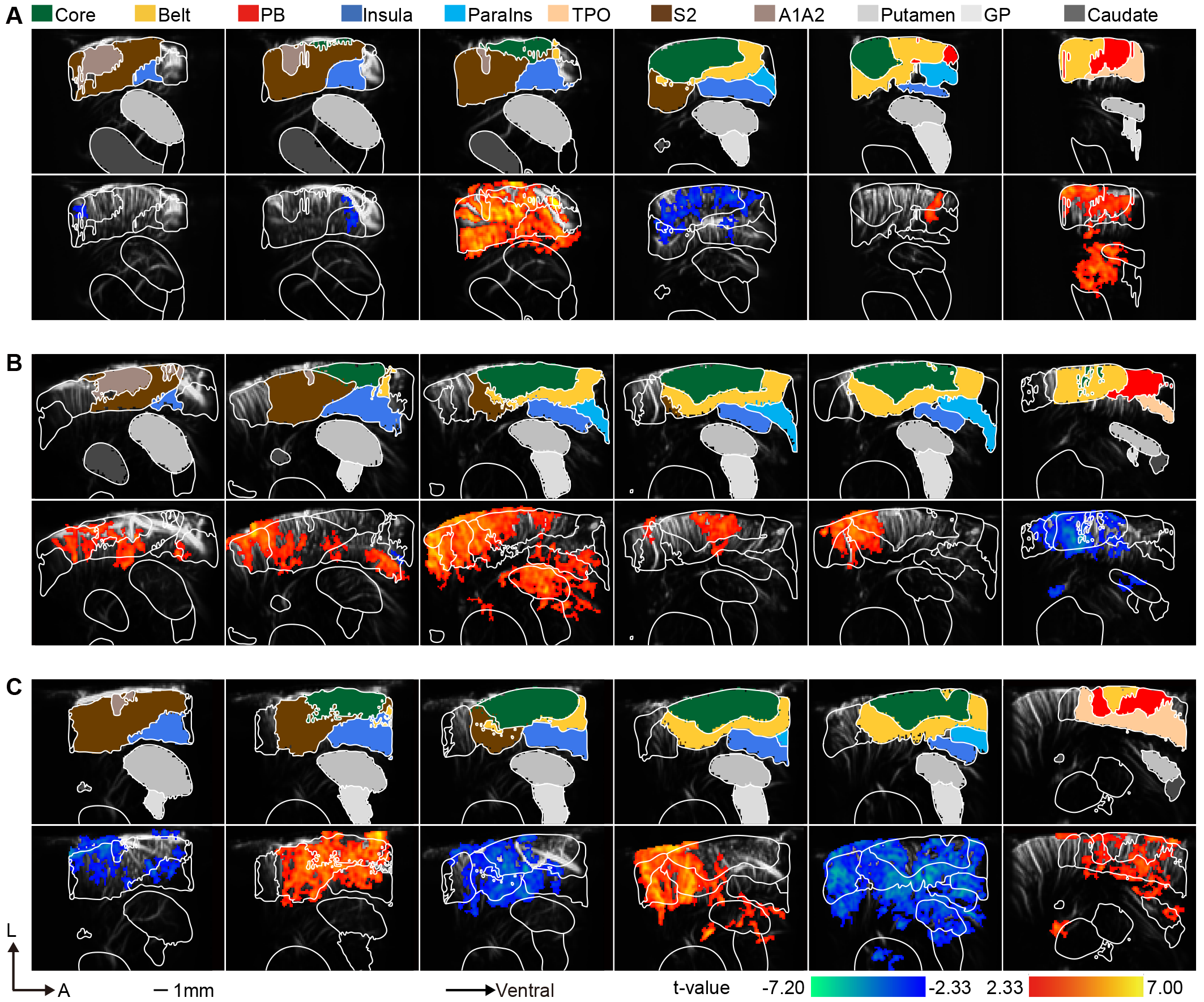

**Figure S7. Individual-level contrast maps evoked by SwapPhrase vs. OriSeries, related to Figure 6**

(A) Top panel: brain region definitions. Bottom panel: contrast map (SwapPhrase vs. OriSeries) for marmoset A2 (GLM, *p* < 0.01, uncorrected).

(B-C) Contrast maps for marmosets A9 (B) and B2 (C) computed using the same analysis settings as in (A).

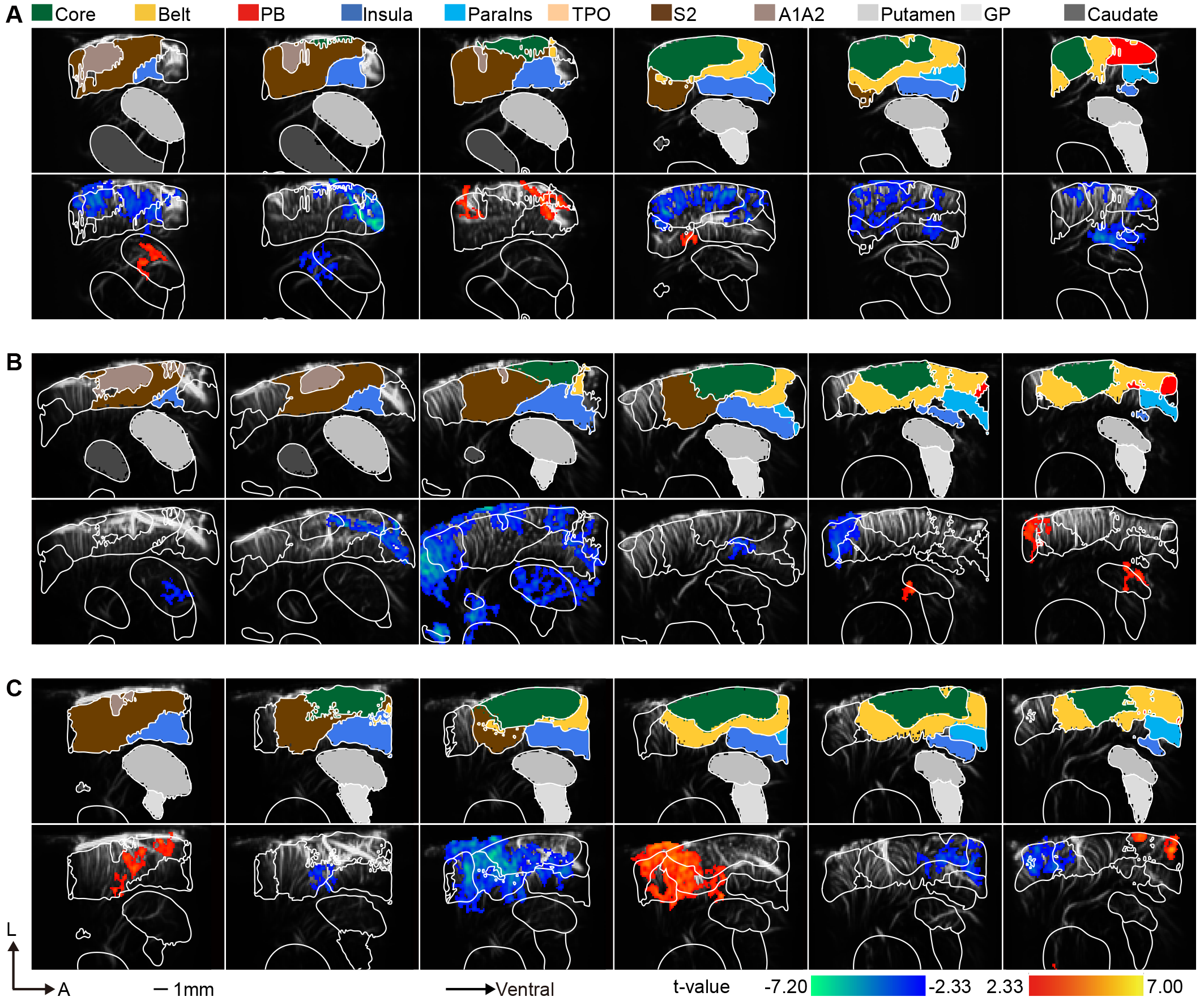

**Figure S8. Individual-level contrast maps evoked by SwapPhee vs. OriSeries, related to Figure 6**

(A) Top panel: brain region definitions. Bottom panel: contrast map (SwapPhee vs. OriSeries) for marmoset A2 (GLM, *p* < 0.01, uncorrected).

(B-C) Contrast maps for marmosets A9 (B) and B2 (C) computed using the same analysis settings as in (A).

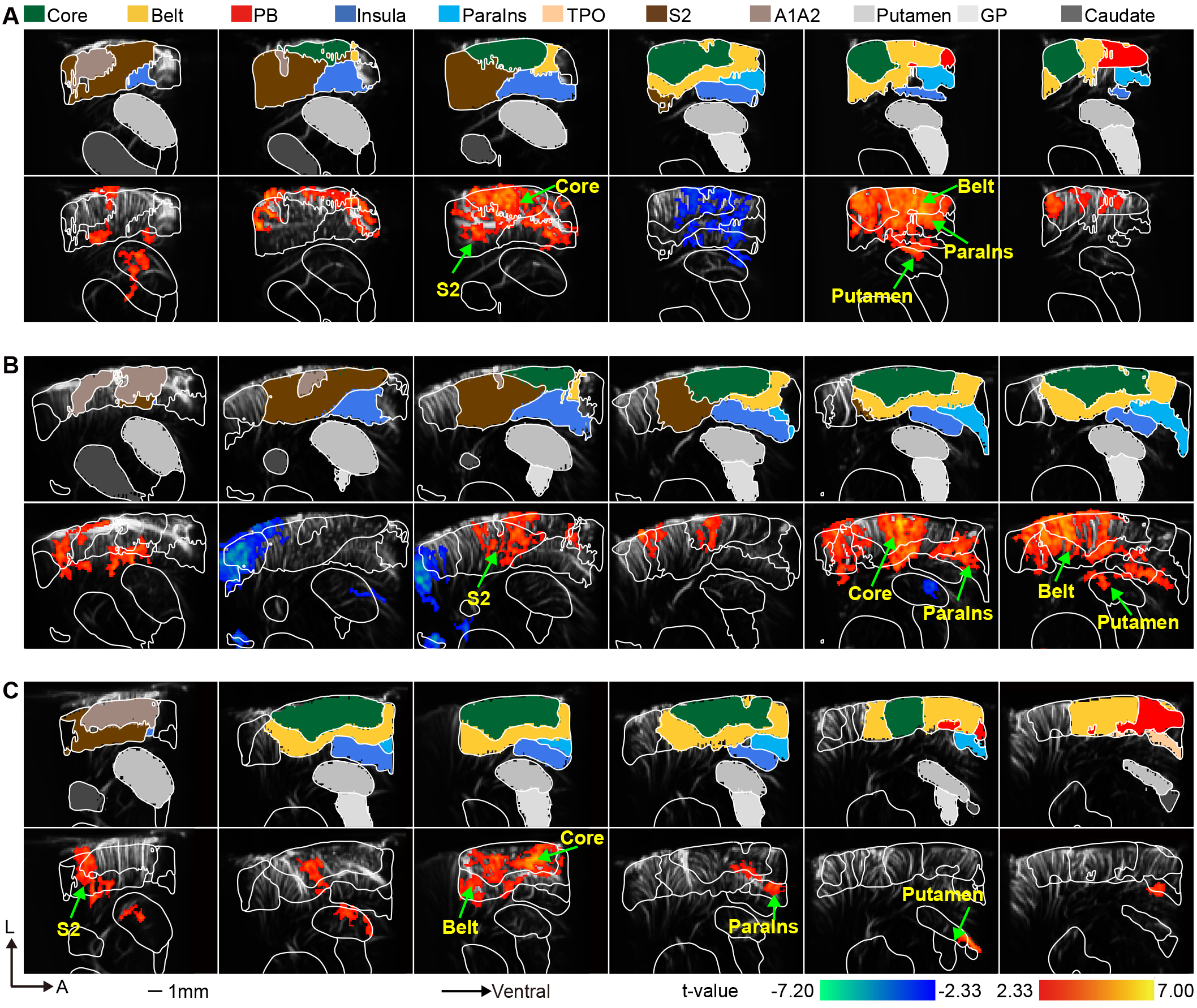

**Figure S9. Individual-level contrast maps evoked by PhraseSpeedUp vs. OriSeries, related to Figure 7**

(A) Top panel: brain region definitions. Bottom panel: contrast map (PhraseSpeedUp vs. OriSeries) for marmoset A2 (GLM, *p* < 0.01, uncorrected).

(B-C) Contrast maps for marmosets A9 (B) and B2 (C) computed using the same analysis settings as in (A).

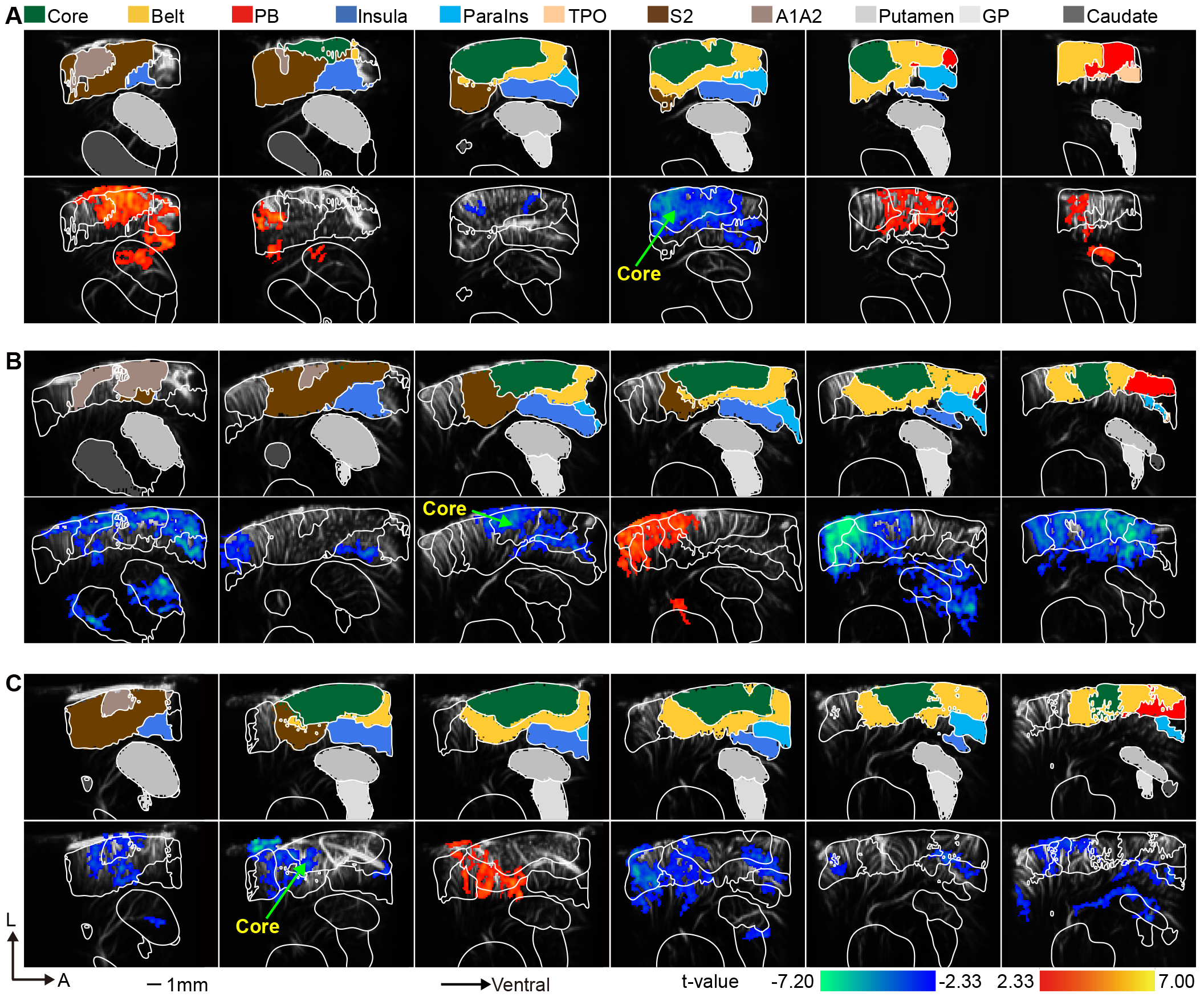

**Figure S10. Individual-level contrast maps evoked by PhraseSpeedDown vs. OriSeries, related to Figure 7**

(A) Top panel: brain region definitions. Bottom panel: contrast map (PhraseSpeedDown vs. OriSeries) for marmoset A2 (GLM, *p* < 0.01, uncorrected).

(B-C) Contrast maps for marmosets A9 (B) and B2 (C) computed using the same analysis settings as in (A).

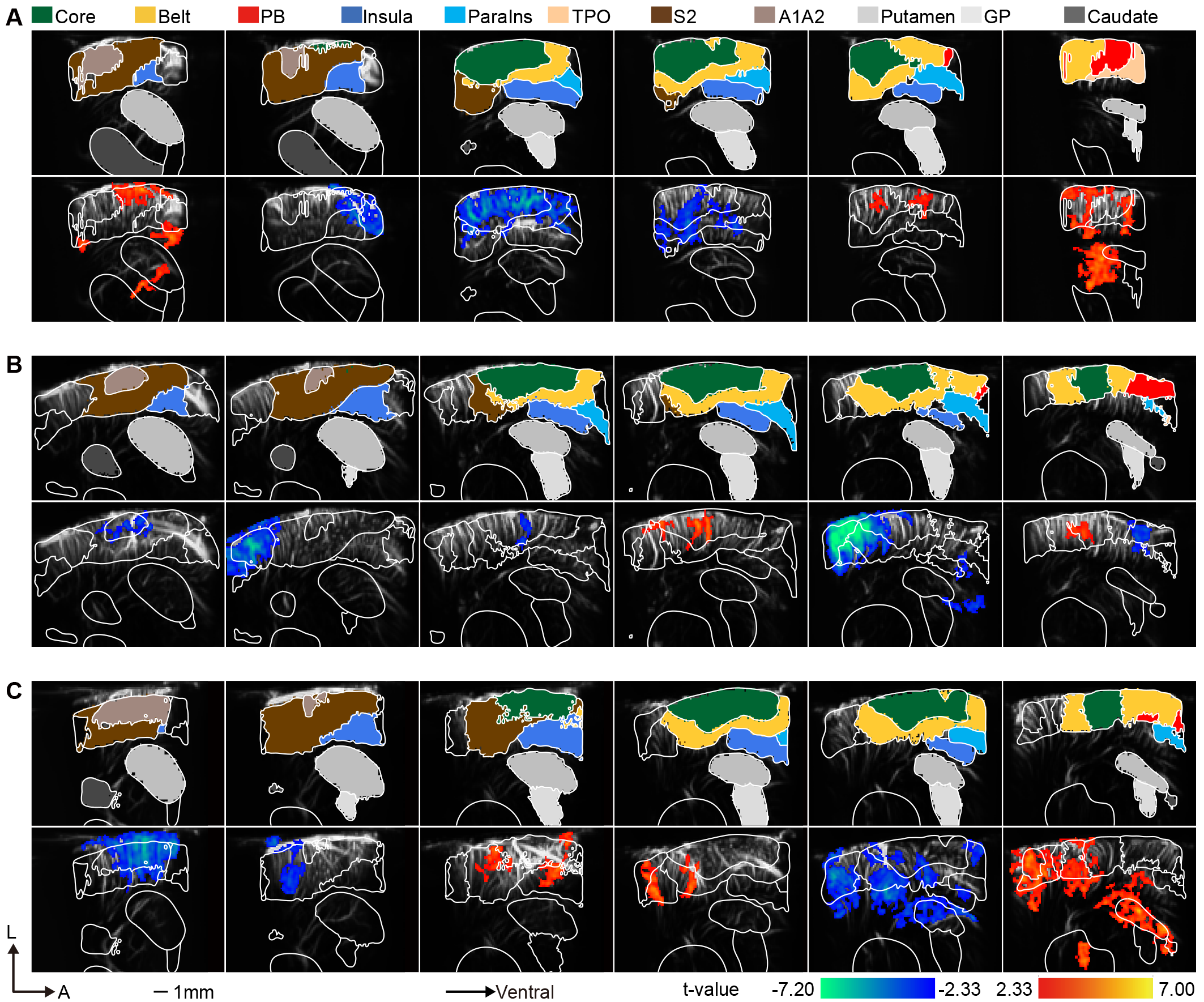

**Figure S11. Individual-level contrast maps evoked by PheeSpeedUp vs. OriSeries, related to Figure 7**

(A) Top panel: brain region definitions. Bottom panel: contrast map (PheeSpeedUp vs. OriSeries) for marmoset A2 (GLM, *p* < 0.01, uncorrected).

(B-C) Contrast maps for marmosets A9 (B) and B2 (C) computed using the same analysis settings as in (A).

**Figure S12. Individual-level contrast maps evoked by PheeSpeedDown vs. OriSeries, related to Figure 7**

(A) Top panel: brain region definitions. Bottom panel: contrast map (PheeSpeedDown vs. OriSeries) for marmoset A2 (GLM, *p* < 0.01, uncorrected; sample size: 33 phee bouts).

(B-C) Contrast maps for marmosets A9 (B) and B2 (C) computed using the same analysis settings as in (A).

**Table 1** Summary of imaging duration and usable lifetime for fUS windows.

| Marmoset ID | Imaging duration (Day) | Usable lifetime (Day) |
| --- | --- | --- |
| A2 | 106 | 237 |
| A9 | 138 | 186 |
| B2 | 260 | 397 |
| B8 | 76 | 129 |

**Table 2** List of ROI abbreviations and full names.

| **Abbreviation** | **Full name** |
| --- | --- |
| AuA1 (Core) | auditory cortex primary area |
| AuR (Core) | auditory cortex rostral area |
| AuRT (Core) | auditory cortex rostrotemporal |
| AuAL (Belt) | auditory cortex anterolateral area |
| AuCL (Belt) | auditory cortex caudolateral area |
| AuCM (Belt) | auditory cortex caudomedial area |
| AuRM (Belt) | auditory cortex rostromedial area |
| AuRTL (Belt) | auditory cortex rostrotemporal lateral area |
| AuRTM (Belt) | auditory cortex rostrotemporal medial area |
| AuRPB (PB) | auditory cortex rostral parabelt |
| Core | auditory core area |
| Belt | auditory belt area |
| PB | auditory parabelt area |
| Insula | insular cortex |
| ParaIns | para-insular cortex |
| TPO | temporo-parieto-occipital association area |
| S2 | secondary somatosensory cortex |
| A1A2 | areas 1 and 2 of cortex |
| GP | globus pallidus |
